# An atlas of the tissue and blood metagenome in cancer reveals novel links between bacteria, viruses and cancer

**DOI:** 10.1101/773200

**Authors:** Sven Borchmann

**Affiliations:** University of Cologne, Department I of Internal Medicine, Center for Integrated Oncology Aachen Bonn Cologne Duesseldorf, German Hodgkin Study Group, Cologne, Germany; University of Cologne, Faculty of Medicine and University Hospital of Cologne, Center for Molecular Medicine, Cologne, Germany; University of Cologne, Faculty of Medicine and University Hospital of Cologne, Else Kröner Forschungskolleg Clonal Evolution in Cancer, Cologne, Germany

## Abstract

Host tissue infections by bacteria and viruses can cause cancer. Massively parallel sequencing now routinely generates datasets large enough to contain detectable traces of bacterial and viral nucleic acids of taxa that colonize the examined tissue or are integrated into the host genome. However, this hidden resource has not been comprehensively studied in large patient cohorts.

In the present study, 3000 whole genome sequencing datasets are leveraged to gain insight into novel links between viruses, bacteria and cancer. The resulting map confirms known links and expands current knowledge by identifying novel associations. Moreover, the detection of certain bacteria or viruses is associated with profound differences in patient and tumor phenotypes, such as patient age, tumor stage, survival, somatic mutations in cancer genes or gene expression profiles.

Overall, these results provide a detailed, unprecedented map of links between viruses, bacteria and cancer that can serve as a reference for future studies.

## INTRODUCTION

Bacterial^1, 2^ and viral^3–5^ infections have widely been recognized as causes of cancer. Examples of carcinogenic viruses are Human Papillomaviridae, causing head and neck^6, 7^ as well as cervical cancer^3, 8, 9^ or Hepatitis B virus, causing liver cancer^10, 11^. The main carcinogenic mechanisms for viral carcinogenesis are thought to be (1) viral integration into and disruption of the host genome and (2) expression of oncogenic viral proteins^12^.

An important example of bacteria causing cancer is Helicobacter which can cause adenocarcinoma of the stomach^13, 14^. The carcinogenic mechanism at play here is thought to be an entirely different one compared to carcinogenic viruses, namely sustained inflammation caused by a chronic, mostly sub-clinical infection^1^. For some links between infections and cancer, preliminary evidence has been presented, but the simultaneous presence of contradictory findings has led to widespread debate. An example of this is the finding that high levels of Fusobacterium nucleatum can be found throughout the cancerous tissue of colorectal cancer at much higher levels than in the tissue of benign adenomas or healthy colon mucosa^15–17^. Given the diversity of carcinogenic mechanisms^18^, it is likely that other carcinogenic viruses and bacteria exist, although currently unknown.

Recent advances in massively parallel sequencing have made it possible to generate large amounts of data informing about the genome, transcriptome and epigenome of a tissue^19^. Resulting datasets contain traces of non-host origin that are present either because of genomic integration or the presence of the virus or bacteria in the tissue itself. While some studies have already been performed with the goal of repurposing this data in order to reveal novel links between infections and cancer^4, 5, 20, 21^, these resources have so far been underutilized.

With the above aim in mind, the present study leverages a large, high-quality collection of over 3000 whole genome sequencing datasets in order to gain insight into novel links between viruses, bacteria and cancer.

## RESULTS

### Samples

A total of 3,025 whole genome sequencing datasets comprising 3.79 trillion reads were included in this study (Figure 1A, Supplementary Table 1). These include 1,330 whole genome sequencing datasets of tumor tissue samples across 14 different cancers from 19 International Cancer Genome Consortium (ICGC)^22^ studies and their corresponding matched normal (e.g. non-cancerous) tissue controls (n=1,330) (Supplementary Table 1). Included patients were predominantly male (n=1,028, 60.6%) and elderly, with 47.3% of patients (n=801) at least 60 years old (Figure 1B-C). Additionally, whole genome sequencing datasets of 365 subjects from the 1,000 genome project^23^ were selected as a healthy control group substituting for the lack of negative sequencing controls to examine non-human DNA in the blood of healthy donors. Only subjects, in whom blood-derived DNA was directly subjected to whole genome sequencing, as opposed to DNA derived from immortalized lymphoblastoid cell lines (LCL) (subset analyzed, n=102), were included in the healthy control cohort. Whole genome sequencing datasets derived from LCL DNA showed a markedly different species-level taxon distribution likely representing taxa present due to LCL culture and not present in the donor itself (Supplementary Figure 2A, Supplementary Data 1-2). For validation purposes and to assess differential gene expression in cancers linked to certain species-level taxa, all available, matching RNA-Seq datasets were also analyzed (n=324).

**Figure 1:**
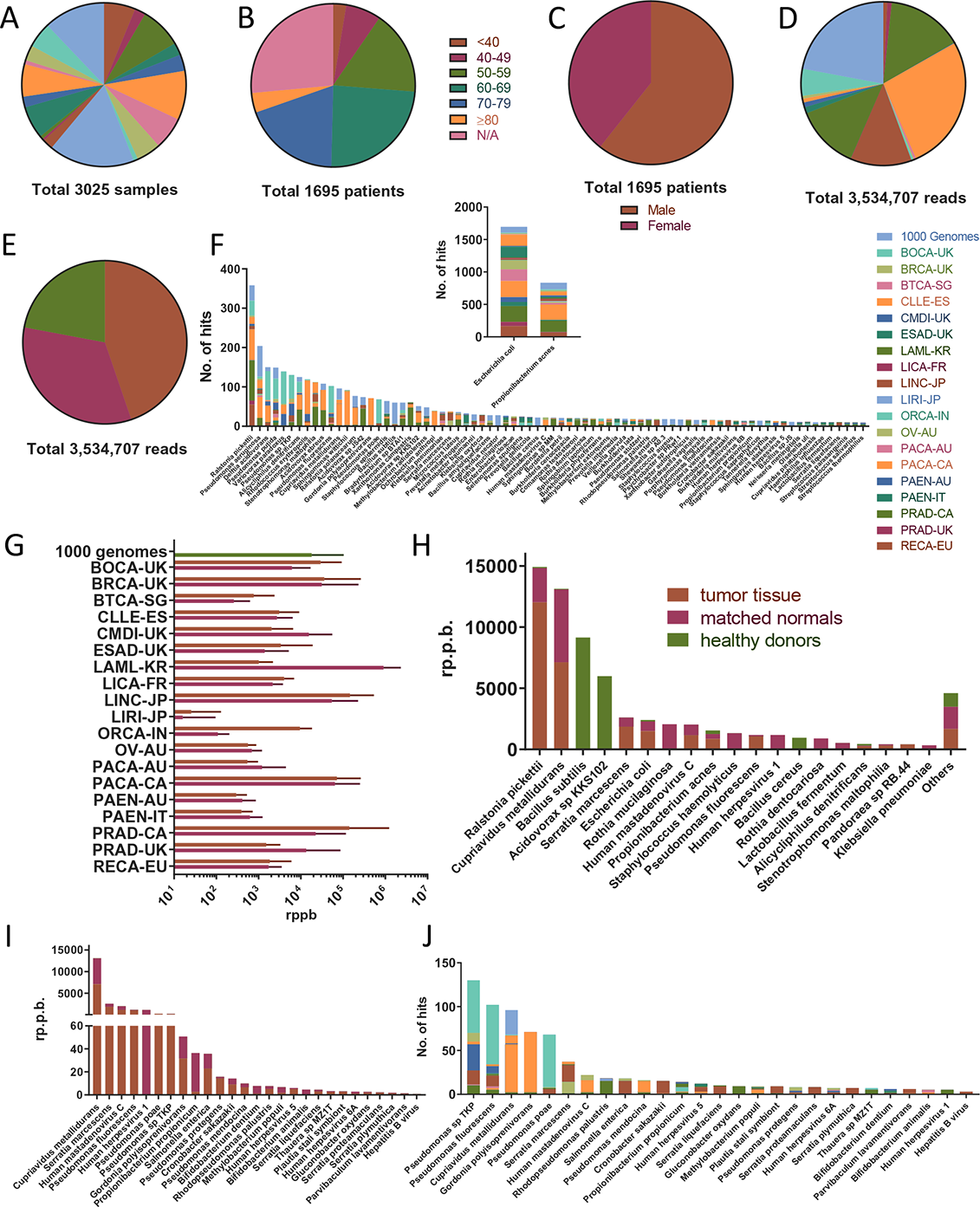
Overview of sample characteristics and identified taxa. A, Sample distribution by project. B, sample distribution by age group. C, sample distribution by gender. D, distribution of read pairs matching any species-level taxa by project. E, distribution of read pairs matching any species-level taxa by sample type. F, species-level taxa detected in at least 10 samples color-coded by project. G, average RPPB detected across all species-level taxa by project and sample type. Bars show mean of samples and error bars show standard error of mean. H, average RPPB detected by species-level taxa and sample type. I, average RPPB detected for all filtered species-level taxa identified as likely tumor-linked by sample type. J, filtered species-level taxa identified as likely tumor-linked color-coded by project.

### Validation of Pipeline

Based on its high accuracy, especially with regard to the precision and high speed necessary for handling large amounts of input data, a pipeline was built around KRAKEN^24^, which at its core is based on the exact alignments of k-mers to their least common ancestor (LCA). Before applying the pipeline to the datasets of interest, it was validated fourfold. In brief, it was confirmed that (i) the pipeline was able to identify already known bacterial and viral taxa in tissue-derived bacterial isolates and cell lines with known integration of viral DNA (Supplementary Data 3, Supplementary Figure 2B), (ii) the taxonomic classification of non-human reads by the pipeline is highly correlated to the classification by an independent approach, namely mapping non-human reads to target genomes with BWA^25^ (Supplementary Data 4, Supplementary Figure 2C-E), (iii) assembled contigs from non-human reads show high sequence similarity with the taxa identified by the pipeline (Supplementary Data 4, Supplementary Figure 2F) and, finally, (iv) identified taxa in pairs of RNA-Seq and WGS data of the same tumor tissue sample are correlated both in a combined dataset of all pairs and within each sample for which RNA-Seq and WGS data was available (n=324) (Supplementary Figure 2 G-H). More details are available in the Supplementary note.

### A map of cancer-linked bacterial and viral taxa

A total of 3,534,707 read pairs matching bacterial, viral or phage species-level taxa were detected across 19 studies in 3025 samples (Figure 1D-E, Supplementary Table 1, Supplementary Data 5). Subsampling 10% of all read pairs did not alter the detected species-level taxa and their relative composition compared to analyzing all non-human read pairs which was validated in a subset of patients (Supplementary Figure 2I-J, Supplementary Data 6). On average, 2.2 species-level taxa per sample were detected, although variation was high (Supplementary Data 7, Supplementary Figure 3 A-D). A total of 218 species-level taxa could be identified in all examined samples (Figure 1F, Supplementary Data 8). Escherichia coli and Propionibacterium acnes were the most commonly detected species in all samples.

In order to control for differences in sequencing depth between samples, all raw read pair counts were normalized by dividing them by 1,000,000,000 total read pairs. This normalized count was defined as read pairs per billion (RPPB). The mean RPPB detected in healthy control samples from the 1000 genome cohort, matched normal samples and tumor tissue samples were 18,112 (4,238 standard error of mean (s.e.m.)), 20,003(4,295 s.e.m.) and 28,282 (9,081 s.e.m.), respectively, with large variation across samples and sequencing projects (Figure 1G). Of note, particularly high RPPB were detected in matched normal, saliva-derived samples from acute myeloid leukemia (AML) patients, as would be expected from a non-sterile source such as saliva. Ralstonia pickettii was the species with the highest RPPB and almost absent in healthy donors, while being detected at higher levels in tumor tissue compared to matched normal samples (Figure 1H). Except for Bacillus subtilis, Acidovorax sp. KKS102 and Bacillus cereus, all species with very high RPPB were detected predominantly in tumor tissue or matched normal samples. A clustered heatmap of the species-level taxa detected in all samples is provided in Figure 2.

**Figure 2:**
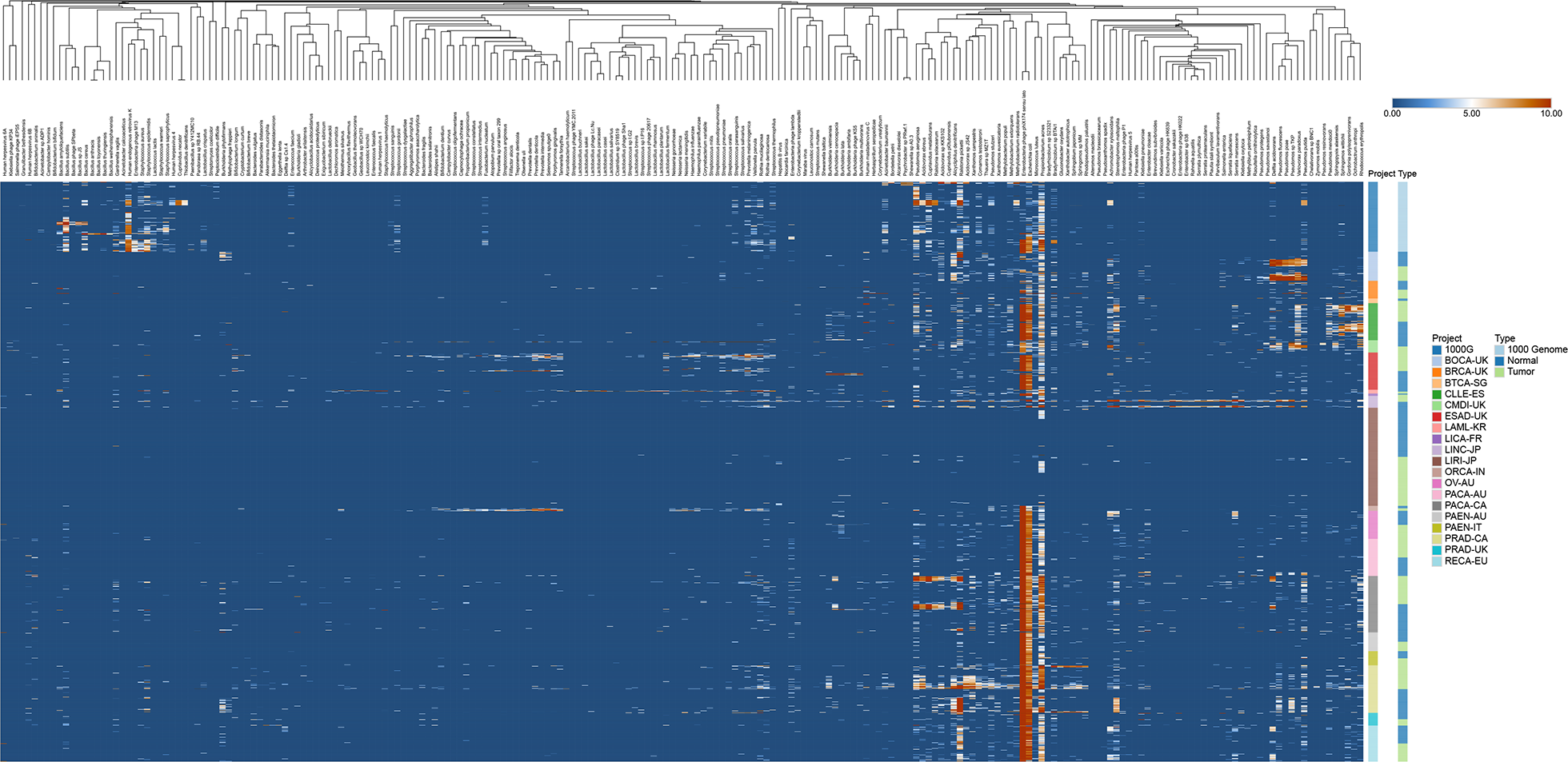
Heatmap of all taxa detected in all samples. Log2-transformed RPPB of all taxa in all samples. Taxa were hierarchically clustered using Pearson correlation as a distance measure with average-linkage. Samples were hierarchically clustered within each project and type subgroup using Pearson correlation as a distance measure with average-linkage.

Next, filtering was performed to exclude taxa that were (i) frequently detected in the healthy control group, (ii) detected in only very few (<5) tumor tissue or matched normal samples, (iii) phages, (iv) taxa that are commonly detected as part of the normal oral microbiome and were mainly detected in saliva, oral or esophageal cancer tissue samples (Supplementary Data 9), (v) taxa that have been previously described as sequencing contaminants (Supplementary Data 10) and (vi) taxa, for which the detected reads were unevenly distributed across the genome (Supplementary Figure 4). After all these filtering steps (Supplementary Figure 5) 27 species-level taxa remained for further analysis (Figure 1 I-J, Supplementary Figure 6, Supplementary Data 11).

Among these, known tumor-linked taxa, such as Hepatitis B virus^10, 11^ or Salmonella enterica^26, 27^ were detected. Furthermore, taxa that have previously been implicated in carcinogenesis, although without enough evidence to support a carcinogenic role, such as Pseudomonas species^28, 29^ and taxa that have never been implicated in carcinogenesis before, such as Gordonia polyisoprenivorans were detected. Of note, most taxa in the final filtered species list were detected at much higher RPPB-levels in tumor tissue compared to matched normal samples. However, lower RPPB of the respective species could still be detected in most matched normal samples. Across all taxa, tumor tissue samples and matched normal samples were highly correlated (Supplementary figure 2K).

In order to better understand the link between detected taxa and different cancers and their relation to each other, a heuristic approach combining the non-linear dimensionality reduction and visualization method t-distributed stochastic neighborhood embedding (t-SNE) with K-means clustering was used. To focus on taxa that potentially play a more direct role in carcinogenesis and considering that RPPB of detected taxa were highly correlated between tumor tissue and matched normal tissue with lower levels detected in matched normal tissue (Supplementary Figure 2K), only tumor tissues were included in the following analyses.

Utilizing a pan-cancer approach, a combined dataset of all tumor tissue samples was visualized using t-SNE using the log_2_-transformed RPPB of all detected non-phage taxa (n=204) as input variables. First, two distinct groups of patients with chronic lymphocytic leukemia (CLL) could be identified. One group (n=24) with detection of Gordonia polyisoprenivorans and another group (n=15) with detection of Pseudomonas mendocina in the tumor tissue. Second, one distinct group (n=23) of pancreatic cancer patients with detection of Cupriavidus metallidurans was observed. Third, one group (n=22) of bone cancer patients with detection of Pseudomonas poae, Pseudomonas fluorescens and Pseudomonas sp. TKP could be distinguished. Fourth, a group of patients (n=14) with bone cancer (n=4) or chronic myeloid disorders (n=10) with detection of Pseudomonas sp. TKP was identified (Figure 3A).

**Figure 3:**
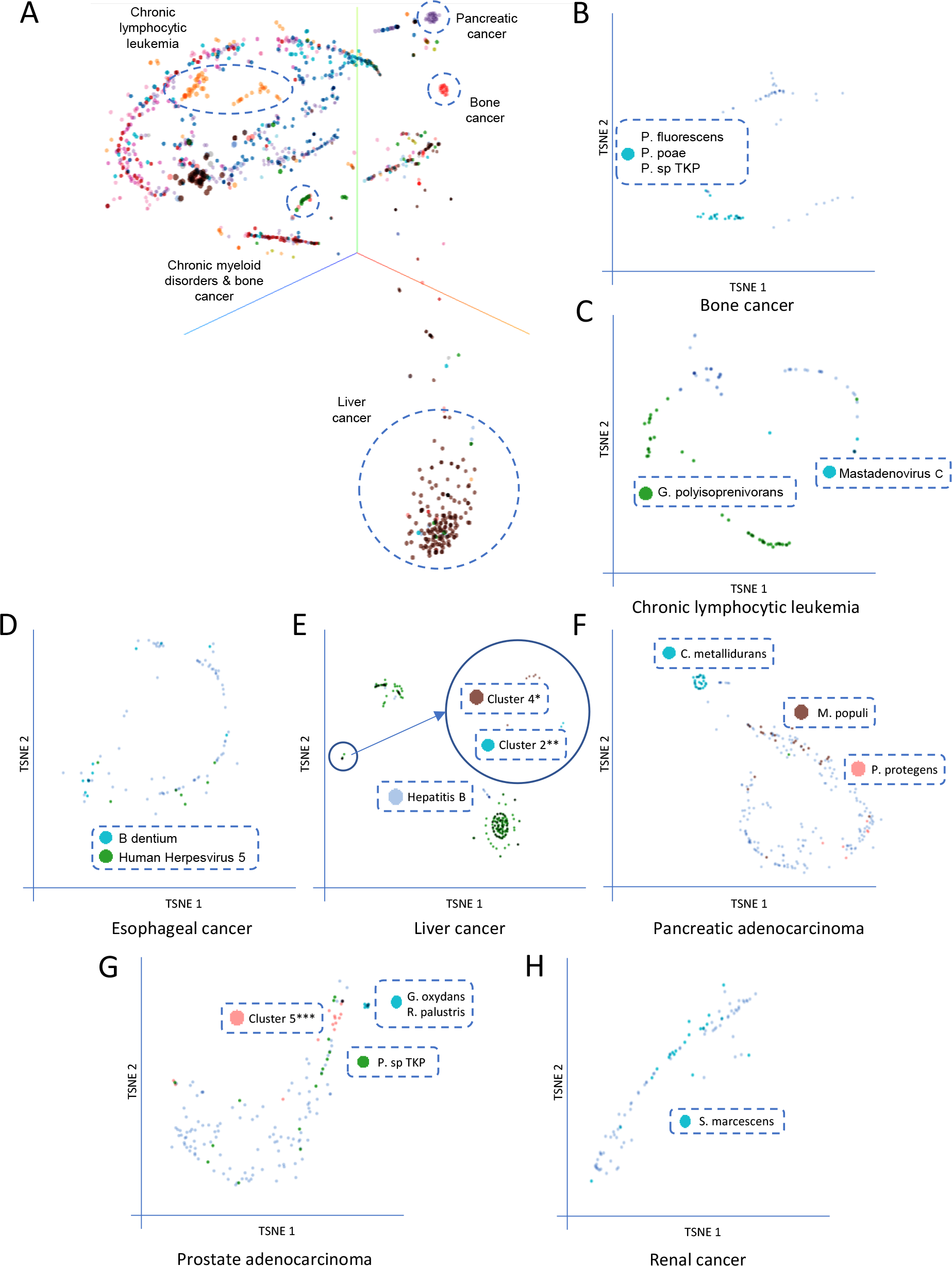
Patient clusters defined by detected taxa can be identified across all patients and in cancer-type subgroups. A, t-SNE visualization of all tumor-tissue samples color coded by project using the log_2_-transformed RPPB of all detected non-phage taxa as input variables. B-H, t-SNE visualizations of single cancer subgroup tumor-tissues color coded by k-means cluster (Supplementary figure 4) using the log_2_-transformed RPPB of all detected non-phage taxa as input variables. *, Pseudomonas sp., Serratia sp. and Salmonella enterica; **, Pseudomonas sp., Serratia sp., Salmonella enterica, Parvibaculum lavamentivorans and Human Herpesvirus 5; ***, Thauera sp. MZ1T, Cupriavidus metallidurans and Pseudomonas mendocina.

In order to gain further insight into links between detected taxa and specific cancers, each cancer-type (Supplementary Data 12) was also analyzed separately. Patient groupings with similar detected taxa were visualized using t-SNE with log_2_-transformed RPPB of all species in the final filtered taxon list as input variables (Supplementary Data 11). Additionally, K-means clustering of patients was performed, using the same input variables. Furthermore, each cluster was linked with age, survival, gender, number of somatic mutations in known cancer genes or specific somatic mutations in one of those cancer genes.

In bone cancer, this dual methodology revealed a cluster of patients with detection of Pseudomonas fluorescens, Pseudomonas sp. TKP and Pseudomonas poae (cluster 2), confirming the results of the pan-cancer analysis (Figure 3B, Figure 4A).

**Figure 4:**
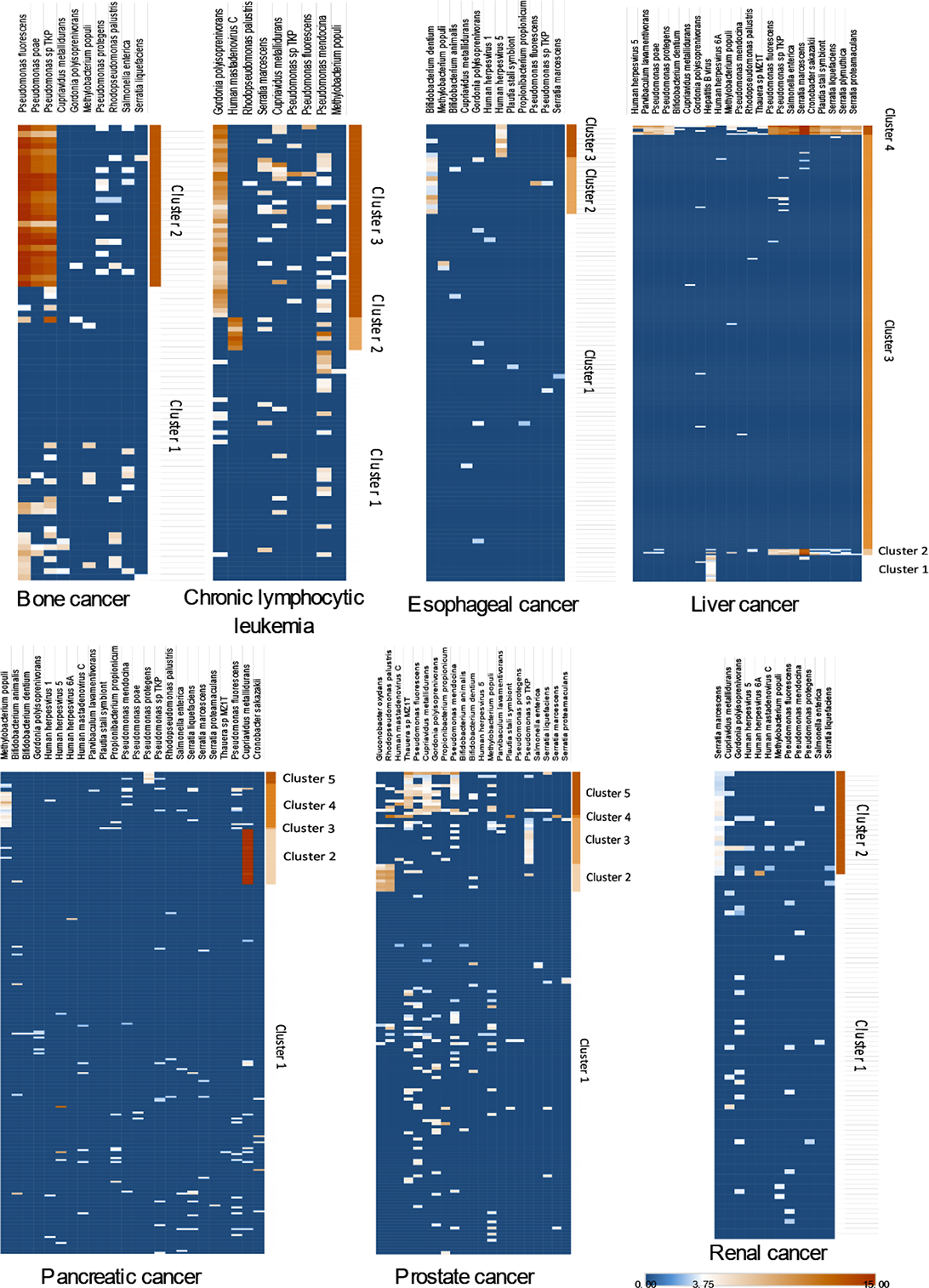
Heatmaps of tumor-linked taxa for all cancers with discernible clusters. A-F, log_2_-transformed RPPB of all species-level taxa identified as likely tumor-linked and detected after filtering in all tumor-tissues of the indicated cancer. Results of K-means clustering of samples are shown.

In chronic lymphocytic leukemia, 2 taxon-linked clusters could be identified, one of patients with detection of Human Mastadenovirus C (cluster 2) and one of patients with detection of Gordonia polyisoprenivorans (cluster 3). Clusters could be identified by both methods, t-SNE and K-means (Figure 3C, Supplementary Figure 3B). Of note, the clusters of patients linked to Human Mastadenovirus C (cluster 2) and Gordonia polyisoprenivorans (cluster 3) were mutually exclusive (p = 0.0124) (Figure 3C, Figure 4B). There was a tendency towards different ages at diagnosis between the clusters (p=0.0743) (Figure 5A), with patients in cluster 2 (Human Mastadenovirus C) being younger. Additionally, there was a tendency towards a difference in survival between the different clusters (p=0.0745). Patients not in any taxon-linked cluster had worse survival than patients in cluster 2 (Human Mastadenovirus C) or 3 (Gordonia polyisoprenivorans) (p=0.0246) (Figure 5B). Patients linked to Gordonia polyisoprenivorans (cluster 3) were more likely to have Binet C stage disease (5/36 vs. 1/61, Binet C vs. not, p=0.0252) (Figure 5C). These patients were also more likely to have TP53 mutations (p(cluster 3 vs. other)=0.0335) (Figure 5D).

**Figure 5:**
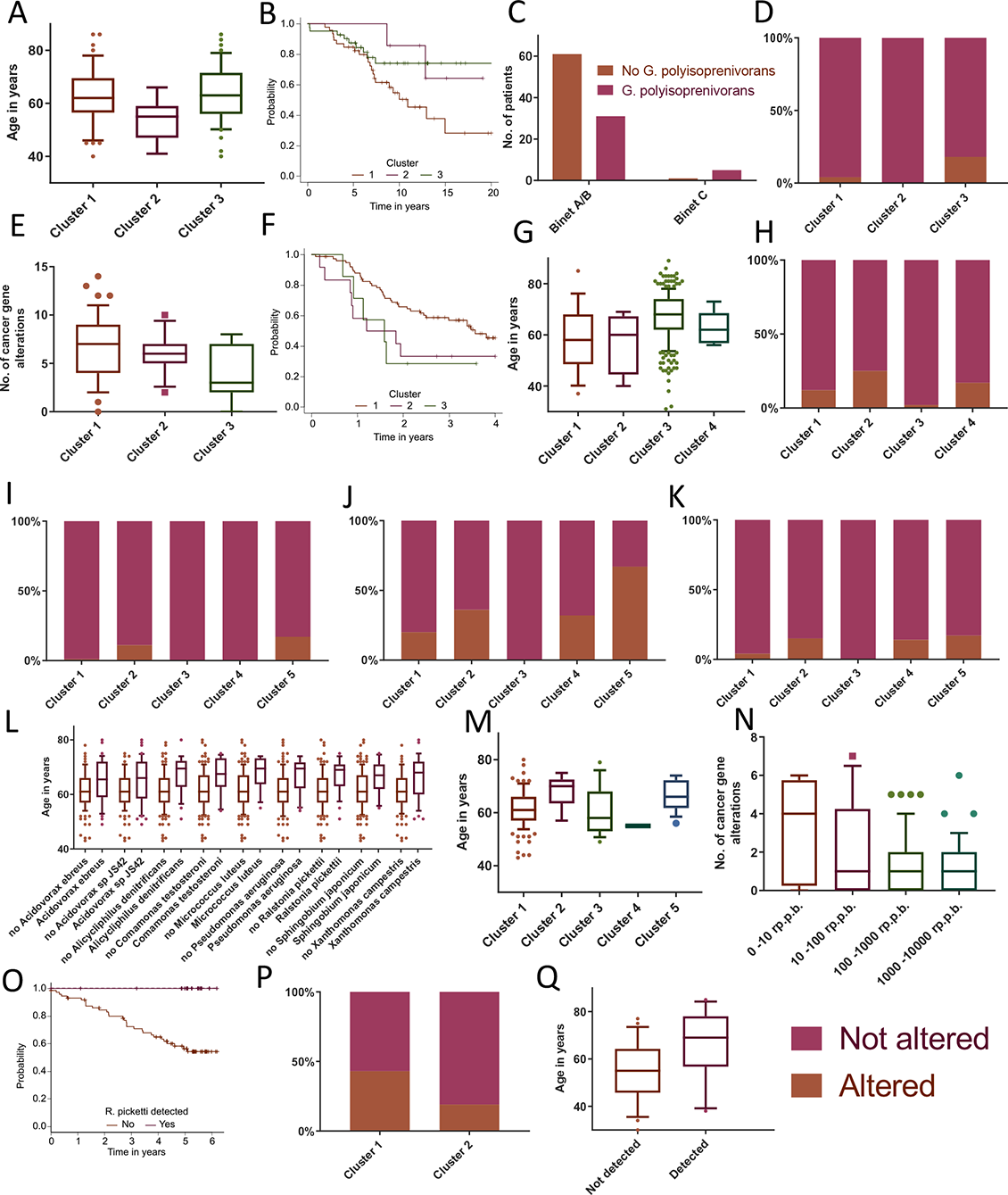
Patient clusters defined by detected taxa are phenotypically distinct. A, CLL patient clusters (1: no specific taxon-link, 2: Human Mastadenovirus C, 3: Gordonia polyisoprenivorans) and age (p=0.0743). B, survival by cluster in CLL (p(1 vs. other)=0.0246). C, Binet stage by detection of Gordonia polyisoprenivorans (p=0.0252). D, TP53 mutation frequency by cluster in CLL (p(cluster 3 vs. other)=0.0335). E, number of cancer consensus gene mutations by cluster (1: no specific taxon-link, 2: Bifidobacterium dentium, 3: Human Herpesvirus 5) in esophageal cancer (p=0.0858). F, Kaplan-Meier survival curves for each cluster in esophageal cancer (p(1 vs. other)= 0.0409). G, liver cancer patient clusters (1: Hepatitis B virus, 2: Pseudomonas sp., Serratia sp. and Salmonella enterica, 3: no specific taxon-link, 4: Parvibaculum lavamentivorans and Human Herpesvirus 5 in addition to taxa from cluster 2) and age (p=0.0015). H, RNF21 mutation frequency by cluster in liver cancer (p=0.0121). I, KMT2C mutation frequency by cluster (1: no specific taxon-link, 2: Cupriavidus metallidurans, 3: no specific taxon-link, 4: Methylobacterium populi, 5: Pseudomonas protegens) in pancreatic cancer (p=0.0308). J, CDKN2A mutation frequency by cluster in pancreatic cancer (p=0.0124). K, RNF21 mutation frequency by cluster in pancreatic cancer (p(RNF21)=0.0107). L, detection of indicated taxa and age in prostate cancer (p_adj_ between 0.0015 and 0.0420). M, prostate cancer patient clusters (1: no specific taxon-link, 2: Gluconobacter oxydans and Rhodopseudomonas palustris, 3: Pseudomonas sp. TKP, 4: no specific taxon-link, 5: Thauera sp. MZ1T, Cupriavidus metallidurans and Pseudomonas mendocina) and age (p=0.0099). N, link between Propionibacterium acne RPPB and number of cancer consensus gene mutations in prostate cancer (p_adj_=0.0041). O, Kaplan-Meier survival analysis of Ralstonia pickettii detection status in renal cancer (p_adj_=0.035). P, PBRM1 mutation frequency by cluster (1: no specific taxon-link, 2: Serratia marcescens) in renal cancer (p=0.0723). Q, Pseudomonas sp. TKP detection and age in chronic myeloid dysplasia (p_adj_=0.039). For all: The midline of the boxplots shows the median, the box borders show upper and lower quartiles, the whiskers show 5^th^ and 95^th^ percentiles and the dots outliers.

In esophageal cancer patients, 2 taxon-linked clusters were identified. K-means clustering revealed one cluster of patients with detection of Bifidobacterium dentium (cluster 2) and one cluster with detection of Human Herpesvirus 5 (cluster 3). However, these 2 clusters could not be differentiated by t-SNE. (Figure 3D, Figure 4C). There was a tendency towards different numbers of somatic mutations in cancer genes between clusters (p=0.0858), with patients in cluster 3 (Human Herpesvirus 5) having fewer mutations than other patients (p=0.0392) (Figure 5E). Additionally, there was a tendency towards a difference in survival between the different clusters (p=0.1). Patients in cluster 2 (Bifidobacterium dentium) or 3 (Human Herpesvirus 5) had worse survival than patients not in any taxon-linked cluster (p=0.040928). (Figure 5F).

Patients with liver cancer could be grouped into 3 taxon-linked clusters by both K-means clustering and t-SNE. One cluster was defined by detection of Hepatitis B (cluster 1). A second cluster was defined by detection of mainly Pseudomonas and Serratia species as well as Salmonella enterica (cluster 2). A third cluster was defined by additional detection of Parvibaculum lavamentivorans and Human Herpesvirus 5 as well as two additional Pseudomonas species, Pseudomonas poae and Pseudomonas protegens (cluster 4), in addition to those detected in the previous cluster. (Figure 3E, Figure 4D). There was a tendency towards different ages at diagnosis between the clusters (p=0.0015) with patients in cluster 1 (Hepatitis B), cluster 2 (Pseudomonas and Serratia sp.) and cluster 4 (Pseudomonas sp., Serratia sp. Parvibaculum lavamentivorans and Human Herpesvirus 5) being younger than patients not in any taxon-linked cluster (p=0.0001) (Figure 5G). In a similar pattern, these patients had a higher frequency of mutations in RNF21 (p=0.0121) (Figure 5H).

In patients with pancreatic adenocarcinoma, 3 taxon-linked clusters could be identified using K-means clustering. One cluster was defined by detection of Cupriavidus metallidurans (cluster 2), one by detection of Methylobacterium populi (cluster 4) and a last one by detection of Pseudomonas protegens (cluster 5). Only the cluster of patients linked to Cupriavidus metallidurans (cluster 2) could also be identified using t-SNE (Figure 3F, Figure 4E). Differences in the frequency of mutations between patients in different clusters were observed in KMT2C, CDKN2A and RNF21. Patients in cluster 2 (Cupriavidus metallidurans), 4 (Methylobacterium populi) or 5 (Pseudomonas protegens) had a higher frequency of mutats in these genes compared to the majority of patients not in any taxon-linked cluster (n=187) (p (KMT2C)=0.0308, p (CDKN2A)=0.0124, p (RNF21)=0.0107) (Figure 5I-K).

In patients with prostate cancer, 3 taxon-linked clusters emerged using K-means clustering, one defined by detection of Gluconobacter oxydans and Rhodopseudomonas palustris (cluster 2), one by detection of Pseudomonas sp. TKP (cluster 3) and a last one by detection of Thauera sp. MZ1T, Cupriavidus metallidurans and Pseudomonas mendocina (cluster 5). Out of these, the cluster defined by Gluconobacter oxydans and Rhodopseudomonas palustris (cluster 2) and the one defined by Thauera sp. MZ1T, Cupriavidus metallidurans and Pseudomonas mendocina (cluster 5) could be confirmed using t-SNE (Figure 3G, Figure 4F). There were age differences at diagnosis observable between the clusters (p=0.0099) with patients in cluster 2 (Gluconobacter oxydans) and 5 (Rhodopseudomonas palustris) being older than the other patients (p=0.0005) (Figure 5M).

Inr patients with renal cancer, a cluster of patients linked to Serratia marcescens (cluster 2) was identified using K-means clustering, although t-SNE did not separate this group of patients (Figure 3H, Figure 4G). There was a tendency towards a lower frequency of PBRM1 mutations in patients in cluster 2 (Serratia marcescens) (p=0.0723) (Figure 5P).

In patients with pancreatic endocrine neoplasms, ovarian cancer, chronic myeloid disorders and breast cancer, no discernable taxon-linked clusters could be identified (Supplementary figure 7A-D).

### Unbiased linkage analysis between bacterial and viral taxa and patient or cancer phenotypes

In addition to linking the above identified clusters with patient or cancer phenotypes, an unbiased analysis of links between the detection of a species-level taxa and patient or cancer phenotypes, such as age, survival, gender, number of somatic mutations in known cancer genes and specific somatic mutations in one of those cancer genes was performed utilizing both a pan-cancer approach and by analyzing each cancer-type separately. In this analysis, all non-phage taxa detected (n=204) were included and multiple testing correction was performed.

When analyzing cancer-types separately, several links were identified. A group of bacterial taxa was linked to older patients in prostate cancer (Figure 5L). In chronic myeloid dysplasia, detection of Pseudomonas sp. TKP was also linked to older age (p_adj_=0.039) (Figure 5Q).

In the cancer-specific analysis, detection of Ralstonia pickettii was linked to improved survival in renal cancer, in fact no patients died (p_adj_=0.035) (Figure 5O).

In prostate cancer, detection of and increasing Propionibacterium acne RPPB were linked to a decreasing number of cancer gene mutations (p_adj_=0.0041) (Figure 5N).

### Somatic lateral gene transfer

The integration of viral nucleic acids into the human host genome is well recognized as a carcinogenic process. High-level evidence exists for the integration of Hepatitis B^10, 11^, Human Papillomavirus^8, 9^ and Epstein–Barr virus (Human Herpesvirus 4)^30, 31^, which are causally linked to hepatocellular carcinoma, cervical cancer and lymphoma, respectively. Additionally, some evidence for somatic lateral gene transfer from bacteria to cancerous tissue has already been presented^20^ and subsequently controversially discussed.

Aiming to find evidence for bacterial or viral DNA integration into the human host genome in this dataset, a pipeline for this purpose was developed. In brief, read pairs in which one read mapped to the human genome and one read to one of the taxa in the final filtered taxon list (n=27) were counted and then compared to the number of read pairs in which both reads mapped to the respective taxa. The number of divergently mapping read pairs divided by the number of complete pairs mapping that respective taxa was used as a measure for genomic integration and lateral gene transfer. This was done for the tumor tissue datasets and the matched normal datasets including all patients (n=79) for which the pipeline revealed at least 1000 RPPB matching one of the taxa in the final filtered taxon list (Supplementary Data 13). The highest rate of integration was observed for Hepatitis B in tumor tissue (9.62%) (Supplementary Figure 8A, Supplementary Data 14). Across all analyzed taxa, putative integrations were more common in matched normal samples than in tumor tissue samples (p=0.0009, Wilcoxon paired signed rank test) (Supplementary Figure 8B, Supplementary Data 14-15) and for viral taxa compared to bacterial taxa (p=0.0001, Mann-Whitney U-test) (Supplementary Figure 8C, Supplementary Data 14-15). In conclusion, there was no evidence for a general phenomenon of lateral gene transfer for any species in the final filtered taxon list, with the exception of Hepatitis B, for which integration into the cancer genome has been widely described^10, 11^.

### Differential gene expression analysis

Differential gene expression analysis was performed for all taxa and cancer combinations with available tumor tissue RNA-seq data, in which at least 5 patients had upwards of 100 RPPB matching the respective taxon (Supplementary Data 16). Differentially expressed genes (q<0.05) were identified for chronic lymphocytic leukemia patients with or without detection of Gordonia polyisoprenivorans (258 genes) (Figure 6A, Supplementary Data 17), Human Mastadenovirus C (1725 genes) (Figure 6B, Supplementary Data 18) and Pseudomonas aeruginosa (50 genes, Supplementary Data 19), respectively. Furthermore, differentially expressed genes were identified for ovarian cancer patients with or without the detection of Escherichia coli (22 genes) (Supplementary Data 20) and pancreatic adenocarcinoma patients with or without detection of Propionibacterium acnes (3 genes) (Supplementary Data 21).

**Figure 6:**
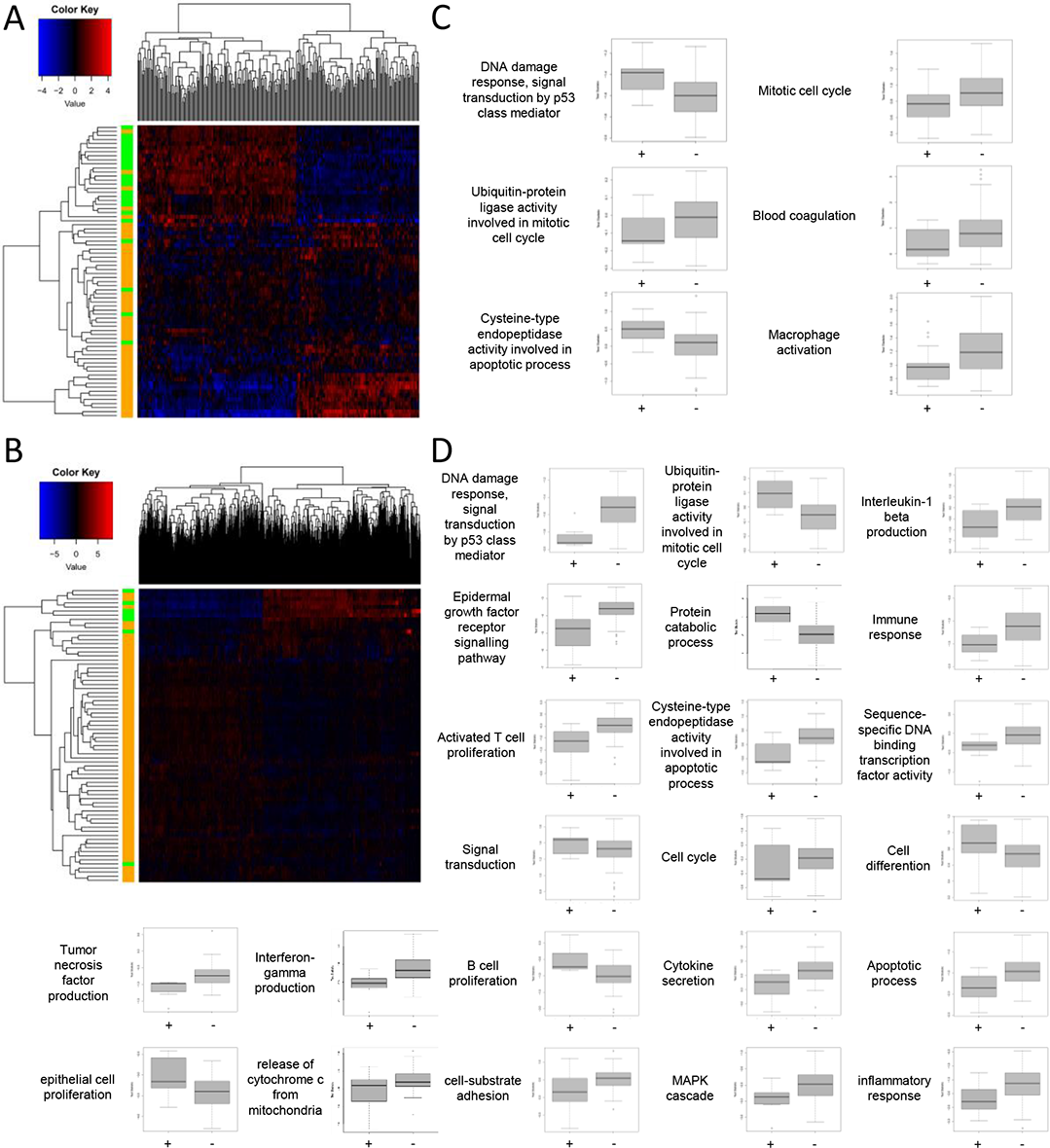
Differential gene expression and pathway analysis of chronic lymphocytic leukemia patients. A, heatmap of all differentially expressed genes (q<0.05) in tumor tissue samples of chronic lymphocytic leukemia (CLL) patients with (green) and without (orange) detection of Gordonia polyisoprenivorans. Dendrograms show clustering with complete linkage and Euclidian distance measure. B, boxplots of the distribution of activating versus inhibitory gene t-test statistics for the indicated pathway is shown for tumor tissue samples of CLL patients with (+, n=23) and without (-, n=49) detection of Gordonia polyisoprenivorans. The midline of the boxplot shows the median, the box borders show upper and lower quartiles and the whiskers the maximum and minimum test statistic. C, heatmap of all differentially expressed genes (q<0.05) in tumor tissue samples of CLL patients with (green) and without (orange) detection of Human Mastadenovirus C. D, boxplots of the distribution of activating versus inhibitory gene t-test statistics for the indicated pathway is shown for tumor tissue samples of CLL patients with (+, n=7) and without (-, n=65) detection of Human Mastadenovirus C. The midline of the boxplot shows the median, the box borders show upper and lower quartiles and the whiskers the maximum and minimum test statistic. Dendrograms show clustering with complete linkage and Euclidian distance measure.

Next, differential gene expression data was used to perform bidirectional functional enrichment in order to identify pathways that are altered in patients with or without detection of the respective taxon. Only the two patient groups with a significant number of differentially expressed genes, chronic lymphocytic leukemia with or without detection of Gordonia polyisoprenivorans (Supplementary Data 22) or Human Mastadenovirus C (Supplementary Data 23) had enriched pathways.

Overall, patients with detection of Gordonia polyisoprenivorans exhibited a gene expression pattern indicative of a decreased level of mitotic cell cycling, an increased DNA damage response, reduced blood coagulation and reduced macrophage activation (Figure 6C).

Contrary to that, patients with detection of Human Mastadenovirus C exhibited a gene expression pattern indicative of an increase in B-cell proliferation, a reduction of Tumor necrosis factor and interleukin beta 1 production, a reduction of activated T cell proliferation and a decrease in cytokine secretion. In addition, the altered tumor tissue gene expression pattern of patients with detection of Human Mastadenovirus C indicated a markedly reduced DNA damage response (Figure 6D).

## DISCUSSION

The aim of this this study was to leverage a large, high-quality dataset of over 3000 samples to reveal novel links between viral and bacterial taxa and cancer. A total of 218 species-level taxa could be identified in tumor tissue, matched normal and healthy donor samples. Out of these, following extensive filtering, 27 taxa were likely cancer-linked. While studies of the viral metagenome of cancer tissues and patients have been performed using datasets from cancer genomics studies^4, 5, 21^, similar large studies examining the bacterial metagenome are lacking.

Studies examining the viral metagenome of cancer tissues mainly identified known links of Human Papillomavirus to cervical and head and neck cancer, of Hepatitis B to liver cancer and of Human Herpesvirus 5 to a variety of cancers while the detection of Human Mastadenovirus C was controversial^4, 5, 21^.

Smaller studies examining bacteria-tumor links in pan-cancer datasets have identified Escherichia coli, Propionibacterium acne and Ralstonia pickettii in multiple cancers, while more specifically finding Acinetobacter sp. in AML and Pseudomonas sp. in both AML and adenocarcinoma of the stomach^20, 32^. Studying cancer-specific datasets, Salmonella enterica, Ralstonia pickettii, Escherichia coli and Pseudomonas sp. were detected in breast cancer and adjacent tissue^33^, while Escherichia sp., Propionibacterium sp., Acinetobacter sp. and Pseudomonas sp. were frequently detected in prostate cancer^29^. Confirming the findings of the present study, these smaller studies found similar bacterial taxa, especially Ralstonia pickettii, Escherichia coli, Propionibacterium acne, Salmonella enterica and Pseudomonas sp. Interestingly, the present study identified a number of taxa that have not been previously identified in cancer tissue or matched normal samples of these patients, among them

Cupriavidus metallidurans, Gordonia polyisoprenivorans, Serratia sp. and Bifidobacterium sp.. Reasons for this are likely (1) the much higher number and diversity of patients and samples included, (2) the larger amounts of data examined per sample due to having higher-coverage WGS datasets available for all patients compared to RNA-seq or whole-exome sequencing (WXS) datasets in previous studies and (3) the optimized and extensively validated bioinformatics approach used here. Of note, the filtering strategy used in this study to exclude taxa likely resulting from contamination has also eliminated bacterial taxa that have previously been linked to cancer such as Escherichia coli and Propionibacterium acne, mainly due to the frequent detection of these taxa in healthy donor samples from the 1000 genome cohort. Stringent filtering for species-level taxa that were detected with at least 100-fold higher RPPB in tumor tissue or matched normal tissue compared to healthy donor samples excluded these species. Despite that, Propionibacterium acne had about 1.4x and 3.1x higher RPPB for matched normal and tumor tissue, respectively, compared to healthy donors. These values were even higher in the case of Escherichia coli, namely 7.1x and 13.4x for matched normal and tumor tissue, respectively. Thus, it is well possible that tumor-linked taxa were eliminated due to the stringent filtering utilized. Therefore, some analyses in this study, such as the unbiased linkage analysis between detected taxa and phenotypes, were performed using the unfiltered dataset and all raw data is provided along with this manuscript for further analysis with different filtering approaches.

Examining specific cancer-pathogen links, a subgroup of bone cancer patients with detection of Pseudomonas sp. in the tumor tissue was identified. Consistent links between viral or bacterial taxa and bone cancer have not been described before, apart from some evidence linking simian virus 40 (SV40) infection to bone cancer^34^. Interestingly, Pseudomonas sp. are frequently implied in difficult to treat cases of osteomyelitis. Thus, one might speculate that chronic, subclinical infection exists and could be carcinogenic.

For CLL patients, two links were identified, one to Gordonia polyisoprenivorans and one to Human Mastadenovirus C. Gordonia polyisoprenivorans has not been linked to cancer before. Interestingly, the bacterium has been identified as a rare cause of bacteremia, so far exclusively in patients with hematological cancers^35–37^. Hematological cancers and their treatment are often associated with profound immunosuppression, allowing for infections with unusual environmental pathogens. It is conceivable that these patients had a latent infection with Gordonia polyisoprenivorans which exacerbated into bacteremia and sepsis upon treatment-induced immunosuppression. To date, Human Mastadenovirus C has not been clearly linked to cancer, although recent reports have found it frequently detected in various cancer tissues, with one treating it as contamination^4^ (see Supplementary Note for a detailed discussion). Further indirect evidence for a role of both Gordonia polyisoprenivorans and Human Mastadenovirus C in CLL carcinogenesis is provided by (1) the observed age difference - patients with detection of Human Mastadenovirus C were markedly younger, (2) the mutual exclusivity of detection of Gordonia polyisoprenivorans and Human Mastadenovirus C and (3) the fact that survival was different - patients with detection of either taxa had improved outcome. This was despite the higher likelihood of patients linked to Gordonia polyisoprenivorans having Binet C stage. Providing an explanation for the observed survival benefit, patients without a taxon-link had a higher likelihood of having TP53 mutations, which are prognostically disadvantageous in CLL^38^. Strikingly, marked differences in host cancer tissue gene expression were observed for patients in which one of the taxa was detected with differences between cases with detection of Gordonia polyisoprenivorans and Mastadenovirus C. The observed tumor tissue gene expression pattern for patients linked to Gordonia polyisoprenivorans, especially an increased DNA damage response and reduced mitotic cell cycling, could explain the improved survival of these patients. The tumor tissue gene expression pattern of patients linked to Human Mastadenovirus C pointed towards reduced immune activity, especially reduced T cell function and a decrease in cytokine production and secretion, providing a potential explanation for the observed high detection frequency of Human Mastadenovirus C DNA in line with an uncontrolled infection due to a diminished immune response. Detection of Mastadenovirus C was linked to a decreased DNA damage response, which has been described as an important pathomechanism of CLL^39^. In addition to CLL patients, both Gordonia polyisoprenivorans and Human Mastadenovirus C were also detected frequently in other cancers, sometimes at high levels, while never being detected in healthy donors.

Two taxon-tumor links in esophageal cancer were identified, one to Bifidobacterium dentium and one to Human Herpesvirus 5. Interestingly, patients with detection of Human Herpesvirus 5 had less somatic mutations in cancer consensus genes than the other patients, pointing to a possibly different carcinogenic mechanism, where the accumulation of multiple somatic mutations is not stringently needed for malignant transformation. Furthermore, patients not in any taxon-linked cluster had better survival. While the link of Human Herpesvirus 5 to esophageal cancer is a novel finding, Human Herpesvirus 5 has frequently been detected in adjacent adenocarcinoma of the stomach^5^. Of note, overt Human Herpesvirus 5 esophagitis can occur in immunocompromised hosts^40^, which could be indicative of a latent infection of the esophagus by Human Herpesvirus 5 in some hosts.

In patients with liver cancer, one subgroup of patients was defined by detection of Hepatitis B, a known cause of liver cancer^10, 11^. Interestingly, two other subgroups could be identified. Pseudomonas sp. and Serratia sp. were detected in both groups, with additional detection of Parvibaculum lavamentivorans and Human Herpesvirus 5 in one group. There is some evidence that Human Herpesvirus 5 might play a role in the carcinogenesis of liver cancer, among them detection of Human Herpesvirus 5 DNA in tumor tissue and an increased seroprevalence in liver cancer patients^41^ as well as frequent hepatitis in Human Herpesvirus 5 infection underscoring hepatotropism of Human Herpesvirus 5^42^. The other identified taxa have not yet been implied in liver cancer carcinogenesis. Interestingly, hepatocellular carcinoma at younger age has been linked to chronic Hepatitis B infection^43^. Similarly, younger age of onset was also observed in liver cancers linked to the other taxa in this study.

Three bacteria-tumor links were identified in pancreatic adenocarcinoma, one with Pseudomonas protegens, one with Methylobacterium populi and one with Cupriavidus metallidurans. While Pseudomonas sp. have been shown to be a contributor to the pancreatic adenocarcinoma tissue microbiome^28^, Methylobacterium populi and Cupriavidus metallidurans have not been detected in pancreatic cancer tissue. In fact, both taxa have not been discovered in human hosts but in environmental samples and are thus not considered part of the human microbiome, making it possible that they are contaminants not truly present in the tumor tissue samples analyzed. On the other hand, few infections of humans by these taxa have been described^44, 45^ and, of note, the first published report of a Cupriavidus metallidurans infection was a case of septicemia in a patient with a pancreatic tumor^45^.

In patients with prostate cancer, the most interesting findings were age differences between patient clusters defined by the detection of different taxa and a negative correlation of Propionibacterium acne detection and number of mutations in cancer consensus genes. Patients with detection of Gluconobacter oxydans and Rhodopseudomonas palustris as well as patients with detection of Thauera sp. MZ1T, Cupriavidus metallidurans and Pseudomonas mendocina were markedly older than the other patients. Gluconobacter sp., Rhodopseudomonas sp., Cupriavidus sp. and Pseudomonas sp. have previously been identified in prostate cancer and normal prostate tissue^29^, but their precise role in prostate disease is entirely unclear. Propionibacterium acne has been implied as a potential carcinogenic bacterium in prostate cancer^46, 47^, possibly by creating a chronic inflammatory microenvironment^48^. In this study, the Propionibacterium acne detection frequency correlated negatively with the number of somatic mutations in cancer consensus genes. This could point to an alternative driver of carcinogenesis by chronic inflammation in the absence of accumulation of many mutations in cancer driver genes.

In renal cancer, a link to Serratia marcescens was identified in a subgroup of patients. While Serratia marcescens has been described as a frequent cause of urinary tract infections, especially in immunocompromised hosts in a nosocomial setting^49^, it has so far not been implicated in cancer. Interestingly, survival of patients with detection of Ralstonia pickettii in their tumor tissue was markedly improved. Ralstonia pickettii is a bacterium that has been filtered out in this study because of frequent detection in healthy donor samples. It could be speculated that, while not causing any overt infection, low level presence of Ralstonia pickettii in the human host is common and improves immunogenicity of renal cancer, thus, improving outcome. It has been shown, that alterations of local and systemic immunity by the host microbiome influence the anticancer immune response^50^, which might be highly relevant for a naturally immunogenic tumor, such as renal cancer^51^.

The intriguing observation of increased somatic bacteria-human lateral gene transfer by Riley et al. ^20^ could not be made in this study. The only taxa for which more integration into the host genome was observed in tumor tissue compared to matched normal was Hepatitis B, for which integration into the genome of cancerous cells has been well recognized^52^.

In conclusion, the present study provides an unprecedented atlas of links of both bacterial and viral taxa to cancer. In addition to confirming known or recently postulated links, several novel links were identified, laying the groundwork for further studies.

## METHODS

### Data sources

Mapped sequencing data for the included ICGC studies was obtained via the ICGC DCC^22, 53^ and downloaded using customized scripts. The use of controlled access ICGC data for this project was approved by the ICGC data access compliance office. Mapped sequencing data for the 1000 genome healthy control samples was obtained from the 1000 genome FTP server^54^ using customized scripts. Sequencing data used for validation was obtained from the European Nucleotide Archive (ENA)^55^.

### Computing environment

Data analysis was performed using a HP Z4 workstation in a Unix environment either using software as mentioned throughout the methods section or customized scripts. Some analyses were performed employing the Galaxy platform^56^.

### Pipeline for taxonomic classification

First, unmapped (non-human) read pairs were extracted from downloaded sequencing data using Samtools (version 1.7)^57^. Bam files were sorted by query name with Picard Tools (version 2.7.1.1)^58^ and converted to FastQ files using Bedtools (version 2.26.0.0)^59^. Subsequently, Trimmomatic (version 0.36.3)^60^ was used to trim reads with sliding window trimming with an average base quality of 20 across 4 bases as cutoff and dropping resulting reads with a residual length < 50. Read pairs, in which one read was dropped according to these rules were dropped altogether. Remaining read pairs were joined with FASTQ joiner (version 2.0.1)^61^ and converted to FASTA files using the built-in function FASTQ to FASTA (version 1.0.0) from

Galaxy^56^. Next, VSearch (version 1.9.7.0)^62^ was used to mask repetitive sequences by replacing them with Ns using standard settings. These masked and joined read pairs were fed into Kraken (version 1.2.3)^24^ using a database of all bacterial and viral genomes in Refseq. The output of each run was filtered with Kraken (version 1.2.3)^24^ setting a confidence threshold of 0.5. A report combining the output of all samples and runs was generated using Kraken (version 1.2.3)^24^. The output was then arranged using customized scripts in R (beginning with version R 3.3.2. and subsequently updated)^63^ to generate the raw metagenome output of each sample. Next, the raw outputs of each sample running the pipeline on a subset of 10% of the cancer tissue and matched normal sample read pairs and on all non-human read pairs of the 1000 genomes samples were filtered by only including species-level taxa and excluding all species-level taxa that were supported by less than 10 read pairs across all samples using R (beginning with version R 3.3.2. and subsequently updated)^63^. Read pairs assigned to Enterobacteria phage phiX174, which is ubiquitously used as a spike-in control in next generation sequencing were omitted from all counts and analyses as an intended contaminant, except for Figure 2, which aims to visualize the full, raw dataset.

To correct for the variation of sequencing depth across samples, matched read pairs per billion read pairs raw sequence (RPPB) were calculated for each sample and each taxon. A taxon was heuristically considered detected, when a respective sample had at least 100 RPPB assigned to that taxon.

### Filtering strategy

First, all taxa that were also highly prevalent in the healthy control group, were excluded from further analysis. In detail, a taxon was required to have a mean RPPB across either the tumor tissue samples or the matched normal samples compared to the healthy control samples of at least 100-fold higher, to be included. This cutoff excluded all taxa that were also highly prevalent in the healthy control samples while at the same time allowing to further analyze taxa that were enriched in matched normal samples such as blood as well as taxa that were dominant in tumor tissue. After this step, 147/218 (67.4%) potential tumor-linked species-level taxa remained. Next, taxa that were detected in fewer than 5 tumor tissue or matched normal samples were excluded (n=78), as well as all remaining phages (n=2). Hepatitis B as a known cancer-linked virus was re-included despite being detected in fewer than 5 tumor tissue or matched normal samples according to these criteria so that 68/218 (31.2%) taxa remained.

Taxa that survived this filtering strategy were likely to be tumor-linked but could also represent artifacts from contamination. To account for that, all taxa were filtered out that are known to be regularly present in the oral microbiome^64–66^ and were at the same time mainly (>50% of all RPPB matching a respective taxon) detected in samples that are likely contaminated with the oral microbiome (saliva matched normal, oral cancer tissue and esophageal cancer tissue samples) (Supplementary data 9). For example, this filtered out taxa that are a commonly present in the oral human flora such as Streptococcus mitis and were indeed mainly detected in tumor tissue samples of oral cancer or esophageal cancer and likely a contaminant based on the biopsy location. Similarly, these taxa were detected in the matched normal of leukemia cases whose matched normal was a saliva sample likely containing taxa of the normal human microbiome. After this filtering step 49/218 (22.5%) species-level taxa remained.

It has recently emerged that both reagents and kits used in DNA extraction and library preparation as well as ultrapure water used in laboratories can contain contaminants that can hamper the detection of truly present taxa in low biomass or high background (e.g. human) samples. As this study was performed using samples that were both low in non-human biomass and in the context of high human background, the aim was to further reduce false positives by compiling a list of common contaminants in microbiome studies. Recommended approaches, such as the sequencing of blank controls^67^ were not feasible as the present study was conducted utilizing already sequenced primary material. Therefore, a database of common contaminants from various studies^68–72^ examining this issue was compiled (Supplementary data 10). All previously recognized contaminant taxa apart from those that were described as a contaminant on genus-level but where different species within the genus were detected differently in 1000 genome control samples and matched normal or tumor tissue samples were excluded. This was the case for the genera Pseudomonas and Methylobacterium. While the species Pseudomonas aeruginosa, Pseudomonas putida and Pseudomonas stutzeri were detected in both 1000 genome control and matched normal or tumor tissue samples and thus likely contaminants, Pseudomonas fluorescens, Pseudomonas mendocina, Pseudomonas poae, Pseudomonas protegens and Pseudomonas sp. TKP were not detected in 1000 genome healthy control samples. Thus, it is likely, that the species-level resolution of this analysis was able to differentiate between common contaminants and possibly tumor-linked taxa. Similarly, differentiation was possible between Methylobacterium radiotolerans and Methylobacterium extorquens (likely contaminants) and Methylobacterium populi, which was only found in tumor samples. The same held true for Cupriavidus metallidurans and Propionibacterium propionicum. After this filtering step, 27/218 (12.4%) species-level taxa remained (Figure 1 I-J, Supplementary Figure 6). While it cannot be ruled out that all these filtering steps removed truly cancer-linked taxa, the aim was to be cautious and rather accept a false-negative than a false-positive finding. Of course, experimental validation of the relevance of one of the taxa eliminated by one of the filters could prove that this taxon is both, a sequencing contaminant and a relevant taxon in cancer.

If a taxon is indeed present in a tissue, matching read pairs are expected to be uniformly distributed across its genome. Consequently, a further filtering step was introduced. The sequencing data from all samples was combined and matched against a reference database constructed out of the genome of these 27 taxa. Next, the coverage distribution of reads across each genome was assessed (Supplementary Figure 4). If detection of the respective taxa is the result from misalignment or sequence similarity between the taxa and for example cloning vectors used in the production of sequencing reagents an uneven coverage would result. It was found that all read pairs matching Human Mastadenovirus C aligned to short parts of its genome with a maximum length of a few hundred base pairs and abrupt drops in coverage (Supplementary Figure 4). This was also found in another study using different cancer tissue sequencing data and resulted in excluding Mastadenovirus C from further analysis^4^. Using Blast (beginning with version 2.7.1 and subsequently updated)^73^ it was found that read pairs matching Human Mastadenovirus C aligned all equally well to commonly used cloning vectors, such as pAxCALGL, which might have been used in the production of reagents used for sequencing. This form of contamination was recently analyzed and found to be frequent^74^. However, read pairs aligning to Human Mastadenovirus C originated from very few, seemingly unlinked samples from diverse cancer sequencing projects, making contamination by recombinant DNA unlikely. Another explanation for the observed coverage pattern is somatic genomic integration of parts of the Human Mastadenovirus C genome into a specific cancer genome. On balance, Human Mastadenovirus C was therefore not excluded from further analysis.

### Clustering of pipeline hits

T-SNE was performed using a web-based TensorFlow Embedding Projector implementation^75^. The learning rate and the perplexity were heuristically set to 10 and 30 for all analyses, respectively, except for the liver cancer subset, for which the perplexity was set to 50 due to improved cluster discrimination. The number of iterations was heuristically chosen, so that no major changes of cluster composition occurred upon increasing the number of iterations. Depending on the subset analysis, between 500 and 1500 iterations were needed to reach that point. T-SNE was performed in 3 dimensions for the pan-cancer analysis and in 2 dimensions for the cancer-specific analyses.

K-means clustering was performed using Morpheus^76^. Unsupervised K-means clustering using Euclidean distance as a similarity measure was employed with the number of clusters being heuristically informed by combining visual inspection, comparing t-SNE and K-means clusters and by examining marginal reduction of within-group variance with increasing numbers of clusters (i.e. the elbow method) (Supplementary Figure 9 A-J).

To combine the information obtained by both complimentary methods, t-SNE clustering was repeated with the same settings for each analysis, while color-coding clusters inferred from K-means clustering.

### Assessment of taxonomic differences between cell culture-derived and blood-derived DNA samples from the 1000 genome project

All taxa in the final taxon list (n=218) (Supplementary data 11) which were identified in blood-derived 1000 genomes healthy control samples were selected (n=76) (Supplementary data 1). The pipeline was subsequently applied to randomly selected (identifier ending with 8 or 9) LCL-derived 1000 genomes samples (n=102) (Supplementary data 2). Finally, RPPB in these LCL-derived samples were calculated for all 76 taxa that were identified in the blood-derived 1000 genomes samples and compared between blood-derived and LCL-derived samples.

### Assessment of effect of subsampling of read pairs on relative taxon distribution

To show that subsampling alters neither the detected species-level taxa nor their relative composition compared to analyzing all non-human read pairs, a subset of 184 tumor tissue samples (Supplementary data 7) was analyzed, without any subsampling and subjected to the pipeline in the same way as in the main analysis. For example, read pairs matching Human Mastadenovirus C, Pseudomonas poae, Ralstonia pickettii and Propionibacterium acnes were used to compare absolute read pair counts between full data and the subsample for all taxon-sample pairs in which the 10% subsample had at least 10 matches to the respective taxon.

### Pipeline validation strategy

In order to perform in-depth validation of the pipeline, two alternative strategies were applied, and results compared.

In the first alternative strategy, BWA mem (version 0.7.17) ^77^ was used to align read pairs to downloaded representative bacterial and viral genomes for the final filtered taxon list (Supplementary data 11). In detail, Samtools (version 1.7)^57^ was used to merge all non-human mapped BAM files. Next, read pairs were extracted using Samtools (version 1.7)^57^ and mapped to phiX (Coliphage phi-X174, complete genome, NC_001422.1) with BWA mem (version 0.7.17)^77^. Using Samtools (version 1.7)^57^, all read pairs not matching phiX were extracted. Picard (version 2.7.1.1)^58^ and Samtools (version 1.7)^57^ were used to extract reads from the resulting BAM file and to sort them by read name. Next, reads were trimmed with Trimmomatic (version 0.36.3)^60^ with sliding window trimming with an average base quality of 20 across 4 bases as cutoff and dropping resulting reads with a residual length < 50. Read pairs, in which one read was dropped according to these rules were dropped altogether. Resulting read pairs were mapped to a merged FASTA file combining reference genomes of the final filtered taxon list (Supplementary data 11). Subsequently, Samtools (version 1.7)^57^ was used to filter only mapped read pairs with a quality of at least 60. Customized scripts were used to generate statistics on the number of mapped reads per species-level taxon in the whole dataset.

In the second alternative strategy, Unicycler (version 0.4.6.0)^78^ was used to assemble reads from a merged read database containing all non-human and non-phiX mapped, trimmed read pairs generated as described above for the first alternative, BWA-based strategy. Unicycler (version 0.4.6.0)^78^ was run with standard settings. The assembled contigs were then matched to sequences in a combined database consisting of the htgs (downloaded on 04/13/2014), nt (downloaded on 04/17/2014 and wgs (downloaded on 04/20/2014) database using BLAST megablast (beginning with version 2.7.1 and subsequently updated)^73^ with an expectation value cutoff of 0.001. Next, customized scripts were used to filter out all contigs with a sequence identity to its matched sequence ≤ 80% and to sum the total alignment length of all remaining contigs by matched sequence template. Finally, for each species in the final filtered taxon list (Supplementary data 11), the matched sequence template representing the species-level taxon with the biggest summed up alignment length was selected.

Next, in order to validate the KRAKEN-based pipeline, the log_10_ of the number of read pairs assigned to a species-level taxon with KRAKEN (version 1.2.3)^24^, the log_10_ of read pairs mapped to a taxon with the BWA-based approach and the log_10_ of the total alignment length of the assembly-based approach were computed for all taxa in the final filtered taxon list (Supplementary data 11) and compared (Supplementary data 4). To assess the correlation between the main method and the two tested alternative methods, the Pearson correlation coefficient was calculated.

Finally, in order to assess the concordance between RNA-Seq and WGS experiments performed on the same sample the main pipeline was applied with the same settings to all samples with RNA-Seq and WGS paired data available (n=324). Pearson correlation coefficients of log_10_ transformed data were calculated for both a combined dataset of all RNA-Seq / WGS pairs and for each sample for which RNA-Seq and WGS data was available (n=324).

### Assessment of somatic lateral gene transfer

In order to assess the integration of bacterial DNA into human DNA, read pairs with one read matching the human genome and one read matching one of the taxa in the final filtered taxon list (n=27) (Supplementary data 11) were identified. First, representative bacterial and viral genomes for the final filtered taxon list were downloaded (Accession numbers in Supplementary data 11). These genomes were merged with the human reference genome (hg1k_v37, downloaded from the 1000 genomes FTP server^54^) into one FASTA file using customized scripts. Second, all read pairs in which only one read of a read pair was mapped to the human genome were filtered using Samtools (version 1.7)^57^. All tumor tissue samples and matched normal samples of all patients in which at least 1000 RPPB matching one of the taxa in the final filtered taxon list were identified in that respective patient’s tumor tissue sample (n=79, Supplementary data 13) and were included in this analysis. BWA mem (version 0.7.17)^77^ with standard settings in paired end mode was used to align all such read pairs to the merged FASTA file of all taxa and the human reference genome. Next, all read pairs that were now divergently mapped were extracted using Samtools (version 1.7)^57^ and customized scripts by only including read pairs where one read mapped to one of the included non-human taxa and the other read mapped to a human sequence. Only read pairs with a mapping quality of at least 40 were retained. Customized scripts were used to count and tabulate all obtained divergently mapped read pairs by taxa and sample, respectively (Supplementary data 14-15).

In order to normalize read pairs mapping to putative integration sites (i.e. divergently mapped read pairs as defined above) by correcting for the total number of read pairs matching a taxon with a similar approach (i.e. non-divergently mapped, putatively non-integrated reads), a comparable pipeline was applied to the data used for the main analysis (i.e. both reads in a read pair not mapped to the human genome) of all patients included in the integration analysis (n=79) (Supplementary data 1). First, the genomes of all taxa in the final filtered taxon list were downloaded (Accession numbers in Supplementary data 11) and merged without adding any further human sequences into one FASTA file to create a reference genome containing all taxa in the final filtered taxon list. BWA mem (version 0.7.17)^77^ with standard settings in paired end mode was used to align these reads to the merged FASTA file of all taxa in the final filtered taxon list. Subsequently, Samtools (version 1.7)^57^ was used to filter the aligned data to only include read pairs that mapped as a proper pair to only one taxon with a minimum mapping quality of 60. Samtools (version 1.7)^57^ and customized scripts were used to count and tabulate all mapped reads. The integration rate of a taxon in either tumor tissue samples or matched normal samples was calculated by dividing the number of divergently mapping read pairs by the number of read pairs mapping as a proper pair to the respective taxon.

### Assessment of links of taxon-defined patient clusters to patient or cancer phenotypes

Differences in age at diagnosis between patient clusters were first analyzed by ANOVA for each cancer. All results with p_anova_≤0.1 are shown in Figure 5. Such clusters or combinations of clusters were then compared to the other clusters by student t-test.

Differences in the gender distribution between patient clusters were analyzed by chi-square test for each cancer.

In order to analyze relationships between the number or type of cancer-associated somatic mutations and detection of specific taxa, all somatic mutations for all patients included in this study were obtained from the ICGC data portal^22, 53^. All synonymous mutations were filtered out. Subsequently, only Tier 1 cancer gene census^79^ genes that were altered in more than 20 cases were filtered using customized scripts and included in the analysis (Supplementary data 24).

Differences in the number of somatic mutations in cancer genes between patient clusters were first analyzed by ANOVA for each cancer All results with p_anova_≤0.1 are shown in Figure 5. Such clusters or combinations of clusters were then compared to the other clusters by student t-test.

The Kaplan-Meier method was used to estimate survival curves for each patient cluster in each cancer-type with available survival data. Differences in survival between patient clusters were analyzed by log rank test.

Links between patient clusters and somatic mutations in single cancer genes were analyzed if the gene was altered in at least 10 patients in that respective cancer-type. Clusters or combinations of clusters were then compared to the other clusters by Fisher’s exact test.

All calculations were performed with R (beginning with version R 3.3.2. and subsequently updated)^63^.

### Unbiased linkage analysis between single bacterial and viral taxa and patient or cancer phenotypes

For this analysis a taxon was considered detected in a patient if at least 100 RPPB matched the respective taxa in the patient’s tumor tissue sample. All phages were excluded from the analysis. All included projects were grouped by cancer (Supplementary data 12). Cancer genes were defined as above.

All calculations were performed with R (beginning with version R 3.3.2. and subsequently updated)^63^. All p-values were corrected for multiple testing using the FDR method to obtain a q-value, which was considered significant if < 0.05.

### Survival analysis

First, the Kaplan-Meier method was used to estimate survival curves for each cancer-taxon pair that was detected in at least 10 patients. Differences in survival between patients with or without detection of a respective taxon were analyzed using log rank tests, stratified by ICGC project. Additionally, a pan-cancer analysis was performed in the same way, also stratifying by ICGC project.

### Links between bacterial or viral taxa and patient gender

Links between detection of a taxa and patient gender were analyzed by Fisher’s exact test for each cancer-taxon pair that was detected in at least 10 patients. Additionally, a pan-cancer analysis was performed in the same way, using a logistic regression model that included the ICGC project as an independent variable.

### Links between bacterial or viral taxa and patient age

Links between the detection of a taxon and patient age at diagnosis were analyzed by student t-test for each cancer-taxon pair that was detected in at least 10 patients. Additionally, a pan-cancer analysis was performed in the same way, using a linear regression model that was stratified by ICGC project.

### Links between bacterial or viral taxa and number of somatic mutations in cancer genes

Links between the detection of a taxon and the number of non-synonymous somatic mutations in cancer consensus genes^79^ of a patient were analyzed by student t-test for each cancer-taxon pair in which a taxon was detected in at least 10 patients. Additionally, a pan-cancer analysis was performed in the same way, using a linear regression model that was stratified by ICGC project.

### Links between bacterial or viral taxa and specific somatic mutations

Links between the detection of a taxon and non-synonymous somatic mutations in one of the cancer consensus genes^79^ of a patient were analyzed by Fisher exact test for each cancer-taxon pair that was detected in at least 10 patients. Additionally, a pan-cancer analysis was performed in the same way, using a logistic regression model that included the ICGC project as an independent variable.

### Assessment of differential gene expression

Reads per kilobase of transcript, per million mapped reads (RPKM) data was downloaded from http://dcc.icgc.org/pcawg for all available tumor tissue samples. Ensemble gene ID was substituted by the standard Human Genome Organization (HUGO) Gene Nomenclature Committee (HGNC) symbol downloaded from http://genenames.org/download/custom and the RPKM data was then linked to the identified species-level taxa in each sample using R (beginning with version R 3.3.2. and subsequently updated)^63^. sRAP^80^ was used to normalize RPKM values, perform quality control, differential gene expression analysis and to identify pathways that are differentially expressed. For these analyses, each cancer was analyzed separately and patients that had more than 100 RPPB matching one species-level taxon in the final filtered taxon list were compared with those who did not. Differential gene expression analysis was performed for all taxa that were detected with more than 100 RPPB in at least five patients and RNA-seq data available (Supplementary data 16). Next, sRAP^80^ was used to perform bidirectional functional enrichment of gene expression data to identify pathways up- or downregulated between patients with or without detection of a taxon. Briefly, the distribution of activation versus inhibition t-test statistics for all samples linked or not linked to a specific taxon was compared using ANOVA and corrected for multiple testing using the FDR method^80^. A gene set was considered functionally enriched if q<0.05. Gene sets likely not relevant for the respective cancer were excluded and gene sets provided with sRAP^80^ were reduced to gene ontology (GO) gene sets^81^.

## SUPPLEMENTARY FIGURES & TABLES

**Supplementary Figure 2:**
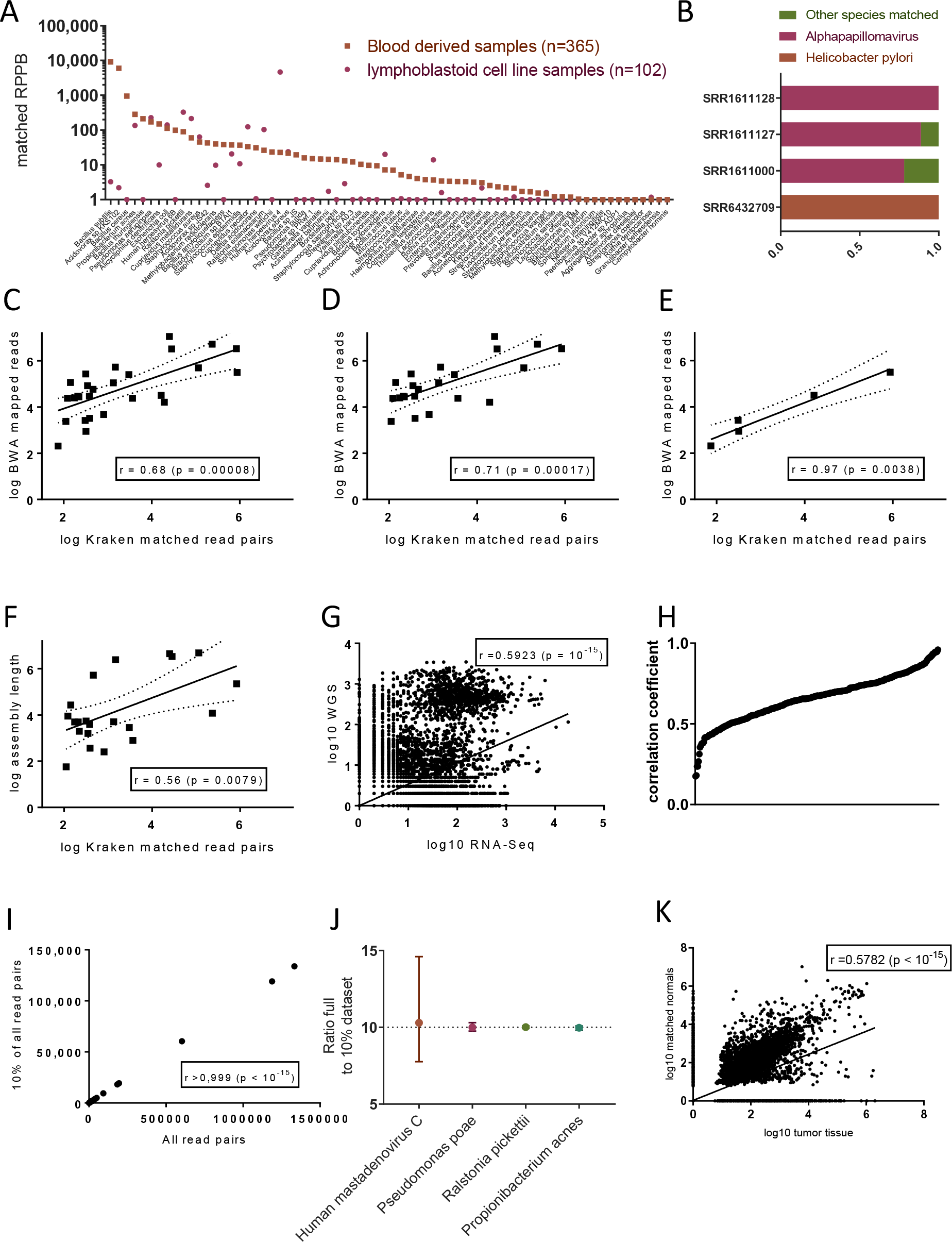
Validation of pipeline and analytical approach. A, mean RPPB detected for indicated species in blood derived and lymphoblastoid cell line 1000 Genome samples sorted by mean of blood derived samples. B, proportion of non-human read pairs matching the indicated taxon of read pairs matching any species-level taxon for each external validation sample. C-E, comparison of BWA mapped reads in Kraken matched read pairs. Each dot represents one species-level taxon (n=27). The line represents the best-fitted line (log-log linear regression). The dotted lines depict the 95% confidence interval. Pearson correlation coefficients (log-log) are shown with two-sided p-values. (C) shows all samples (n=27), (D) only bacterial taxa (n=22) and (E) only viral taxa (n=5). F, comparison of assembled read length and Kraken matched read pairs. Each dot represents one species-level taxon (n=27). The line represents the best-fitted line (log-log linear regression). The dotted lines depict the 95% confidence interval. Pearson correlation coefficients (log-log) are shown with two-sided p-values. G, comparison of Kraken matched read pairs in RNA-seq and WGS data of the same sample. Each dot represents one species-level taxon in one sample with both RNA-seq and WGS data available. The line represents the best-fitted line (log-log linear regression). Pearson correlation coefficients (log-log) are shown with two-sided p-values. H, plot of Pearson correlation coefficient (log-log) distribution of all samples with both RNA-seq and WGS data available. Each dot represents the Pearson correlation coefficient within a single sample. I, comparison of 10% subsample and full dataset. Each dot represents one species-level taxon in one sample for which both the full and the subsampled dataset has been analyzed and indicates the absolute read count identified in both samples. The line represents the best-fitted line (log-log linear regression). Pearson correlation coefficients (log-log) are shown with two-sided p-values. J, Ratio of absolute read counts in the full sample to the 10% subsample for 4 selected taxa. The mean ratio of all samples in which the respective taxon was detected is indicated by the symbol and the error bars indicate the standard error of the mean. The dotted line shows the expected ratio of 10. K, comparison of tumor tissue and matched normal by patient and taxon. Each dot represents one species-level taxon in one patient with both tumor tissue and matched normal analyzed and indicates the RPPB in both samples. The line represents the best-fitted line (log-log linear regression). Pearson correlation coefficients (log-log) are shown with two-sided p-values.

**Supplementary Figure 3:**
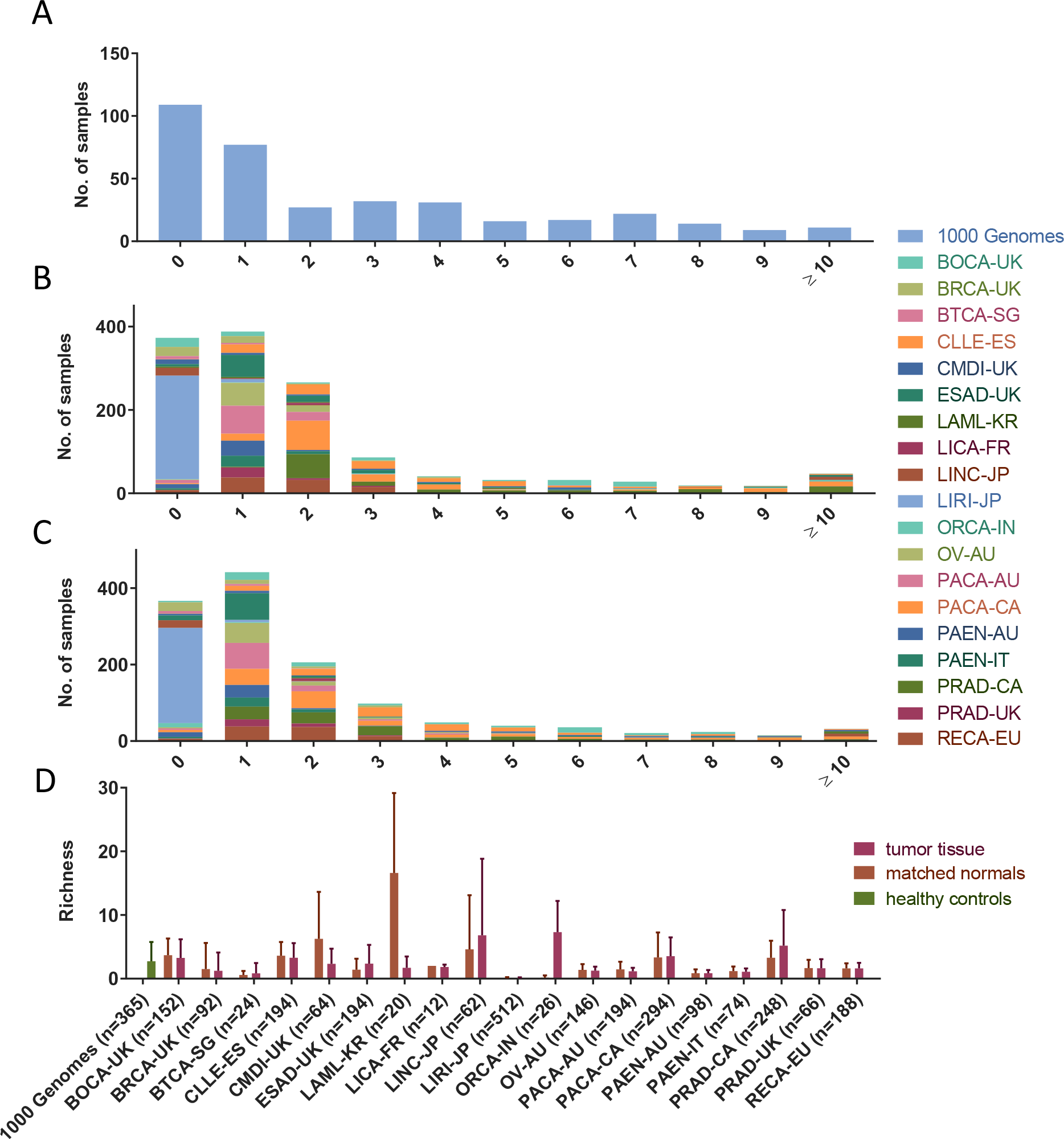
Alpha diversity. A, counts of 1000 Genome samples by species-level richness. B, counts of tumor tissue samples by species-level richness color-coded by project. C, counts of matched normal samples by species-level richness color-coded by project. D, comparison of richness between projects and sample type. Bars show mean and error bars standard deviation. N in brackets indicates total sample number for each project.

**Supplementary Figure 4:**
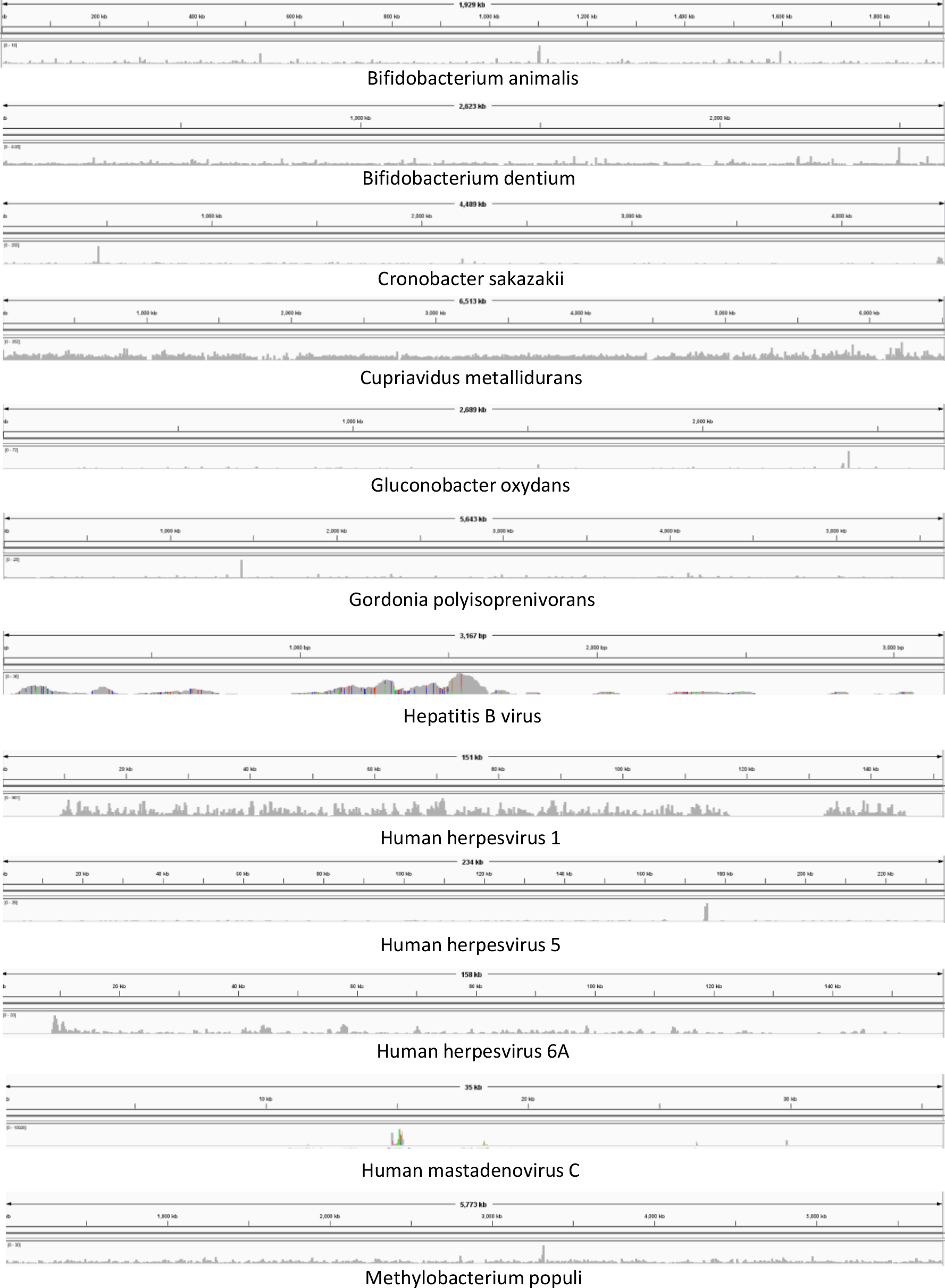
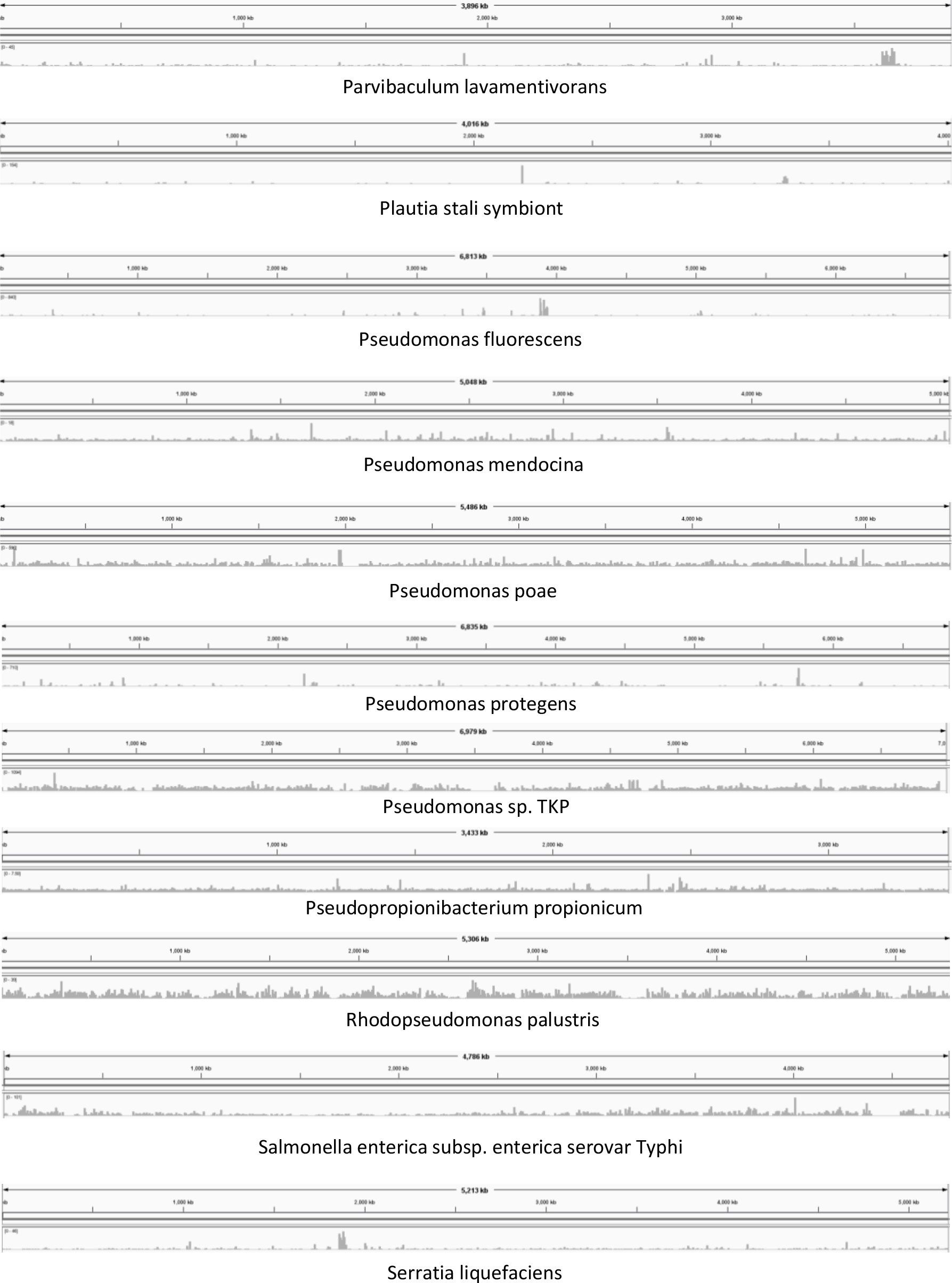
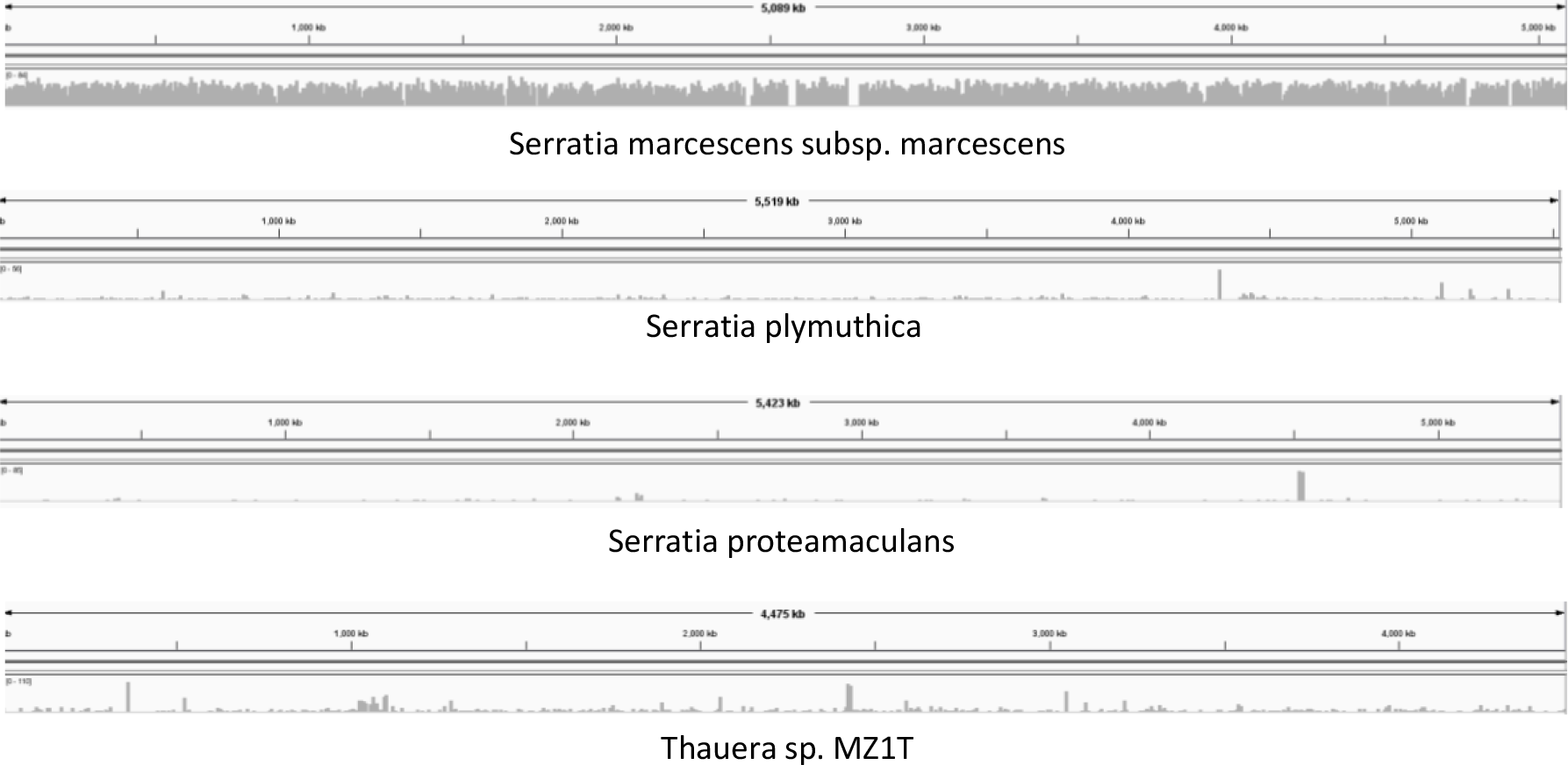
Coverage distribution for all tumor-linked species-level taxa. Coverage distribution across each species-level taxon identified as tumor-linked.

**Supplementary Figure 5:**
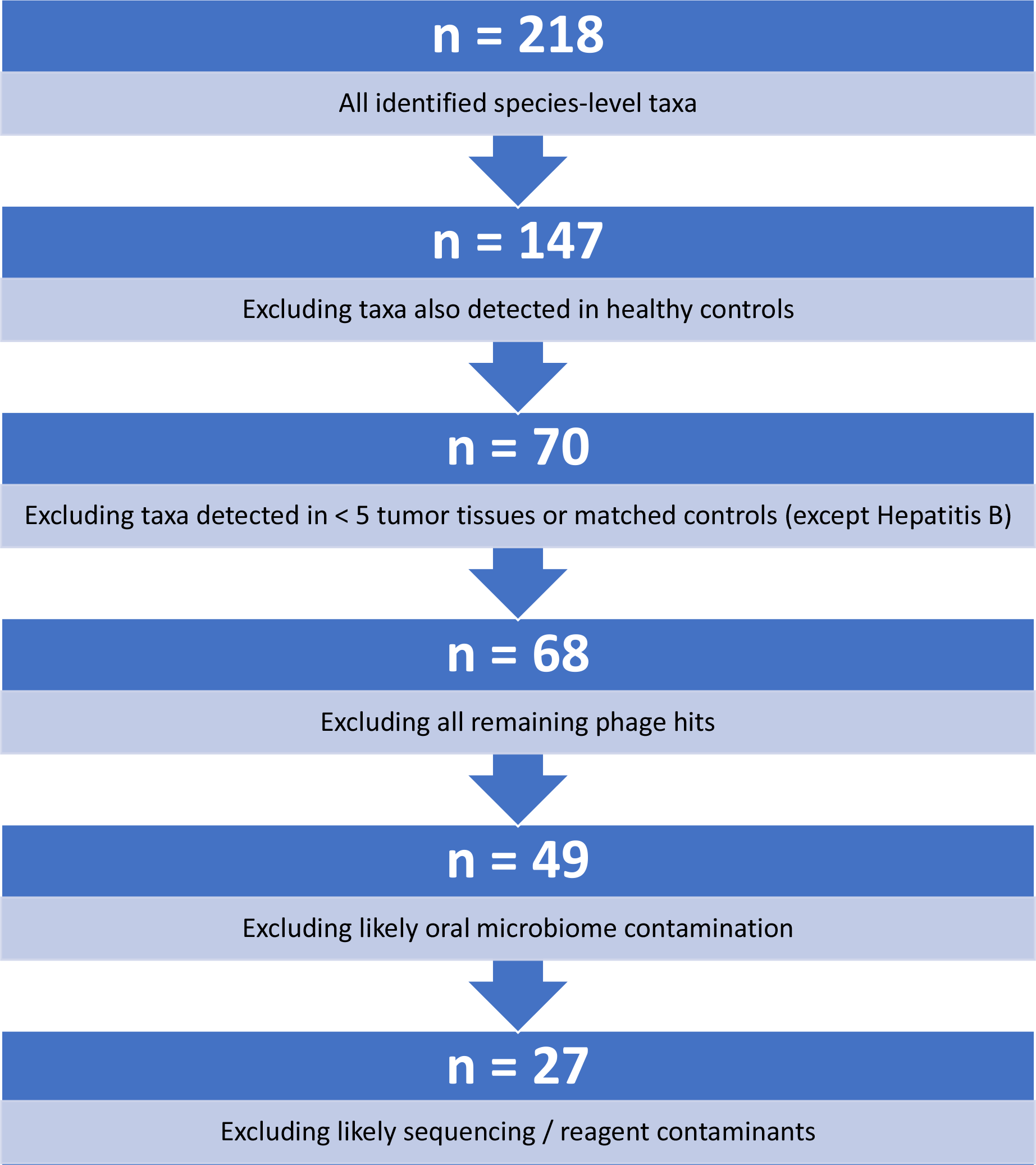
Flow chart of taxa filtering strategy. Flow chart of filtering strategy to derive likely tumor-linked species-level taxa.

**Supplementary Figure 6:**
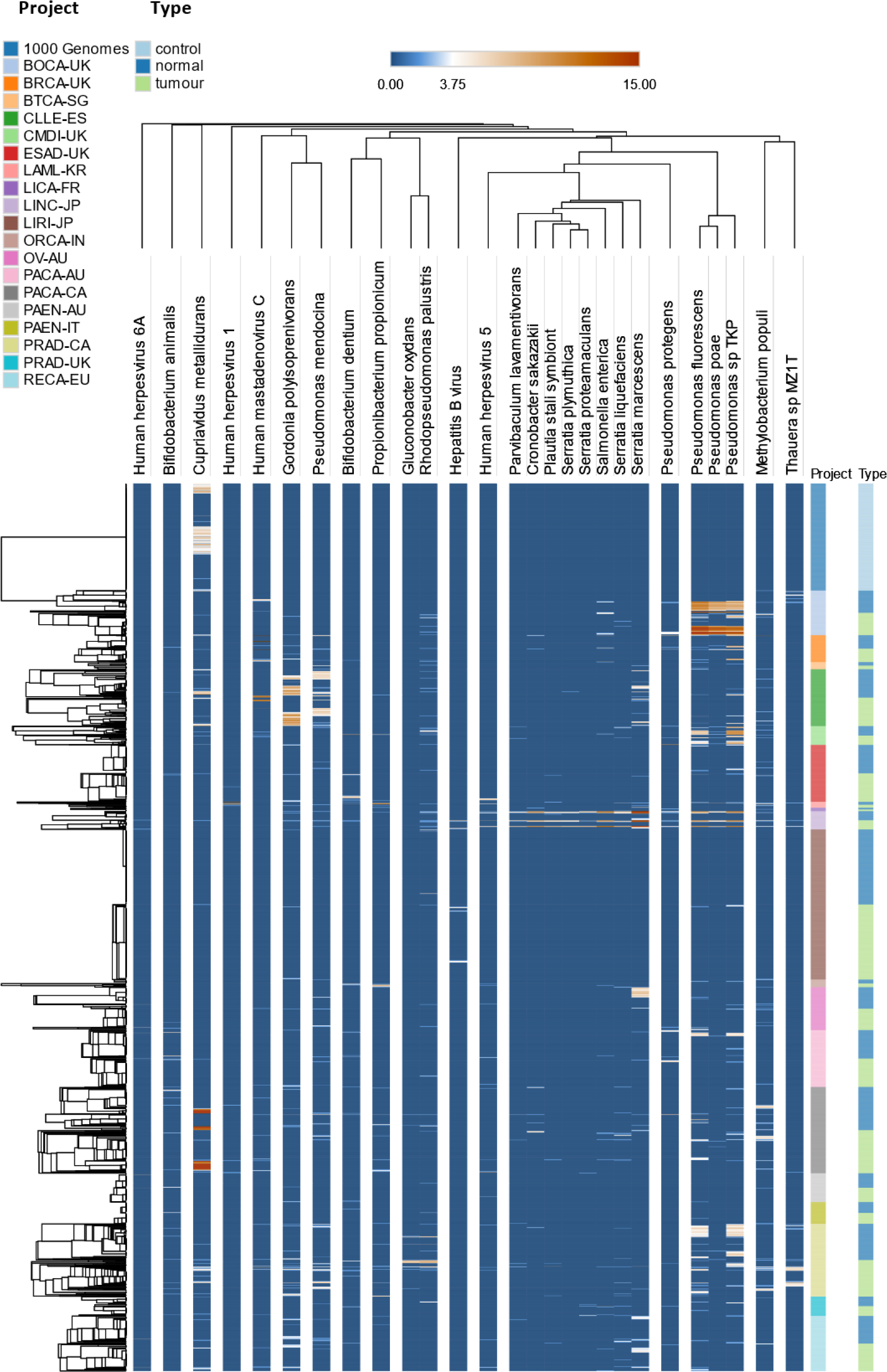
Heatmap of filtered taxons. Log_2_-transformed RPPB of all species-level taxa identified as likely tumor-linked after filtering in all samples. Taxa were hierarchically clustered using Pearson correlation as a distance measure with average-linkage. Samples were hierarchically clustered within each project and type subgroup using Pearson correlation as a distance measure with average-linkage.

**Supplementary Figure 7:**
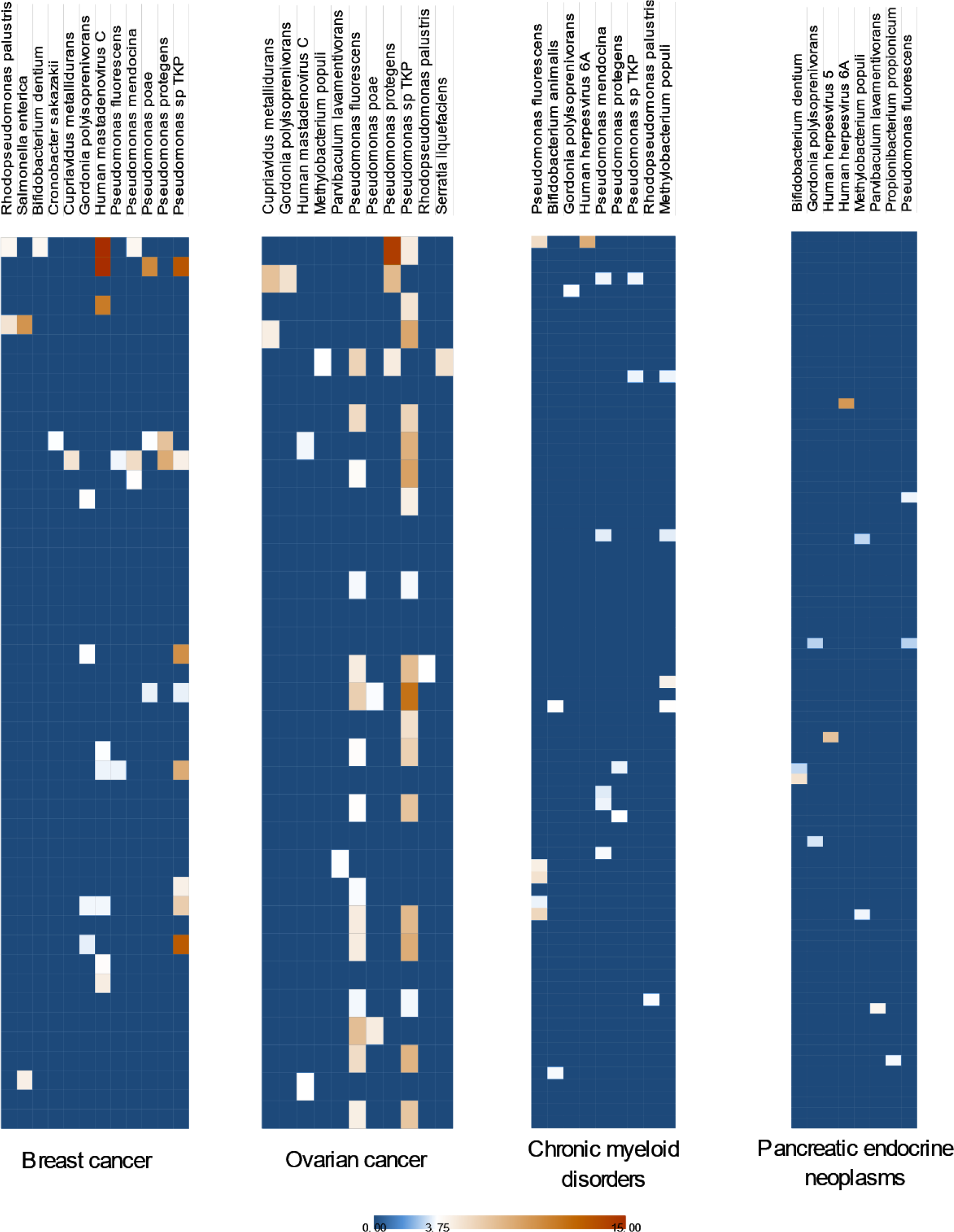
Heatmaps of tumor-linked taxa for all cancers without discernible clusters. A-D, log_2_-transformed RPPB of all species-level taxa identified as tumor-linked and detected after filtering in all tumor-tissues of the indicated cancer. Results of K-means clustering of samples are shown.

**Supplementary Figure 8:**
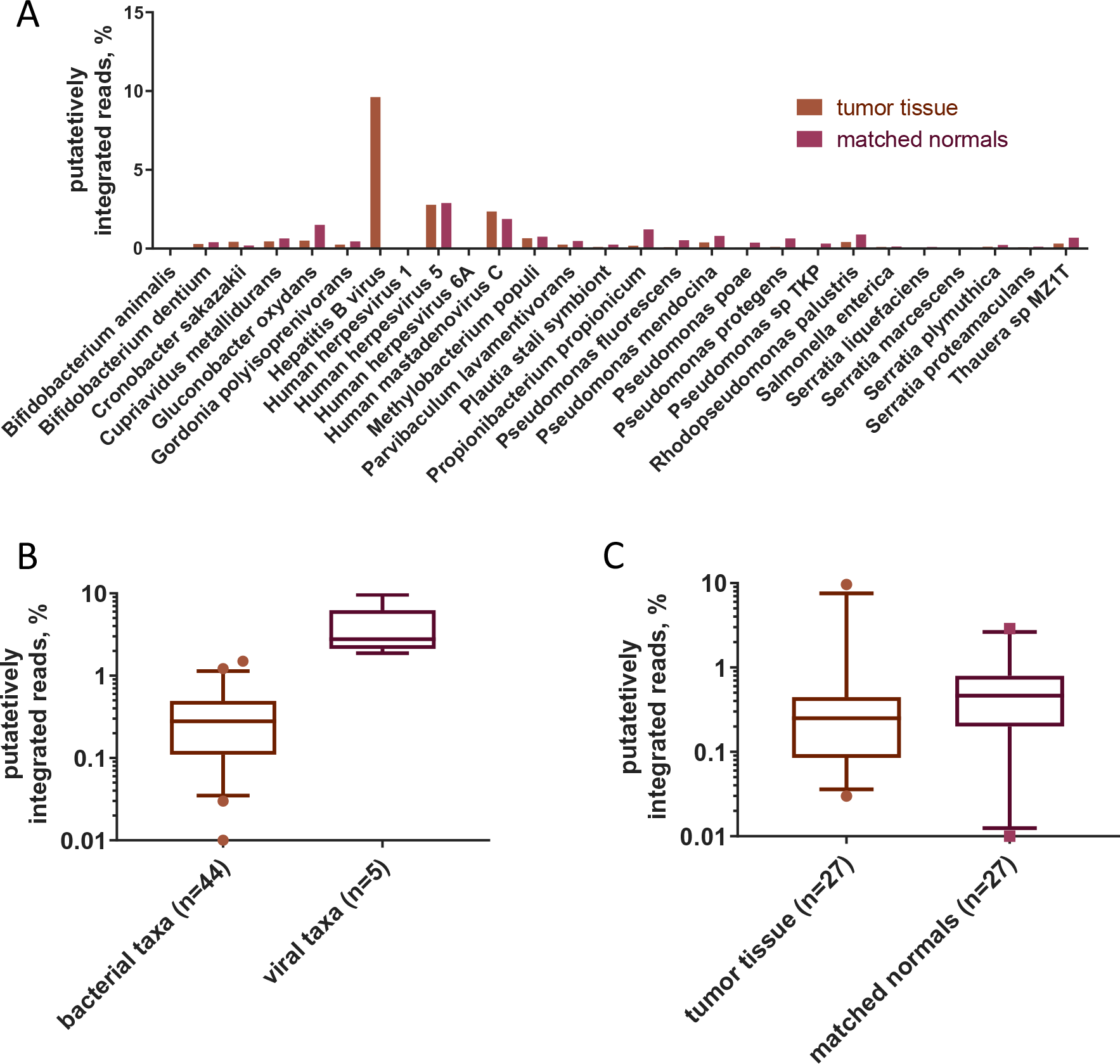
Host integration. A, integration rate by species for tumor tissue and matched normal sample. B, difference in integration rates between bacterial and viral taxa (p<0.0001, Wilcoxon rank-sum test, two-tailed). The midline of the boxplot shows the median, the box borders show upper and lower quartile, the whiskers show 5^th^ and 95^th^ percentiles and the dots outliers of species-specific integration rates in tumor tissue or matched normal samples. C, difference in integration rate between tumor tissue and matched normal samples (p=0.0009, Wilcoxon signed-rank test, two-tailed). The midline of the boxplot shows the median, the box borders show upper and lower quartile, the whiskers show 5^th^ and 95^th^ percentiles and the dots outliers of species-specific integration rates.

**Supplementary Figure 9:**
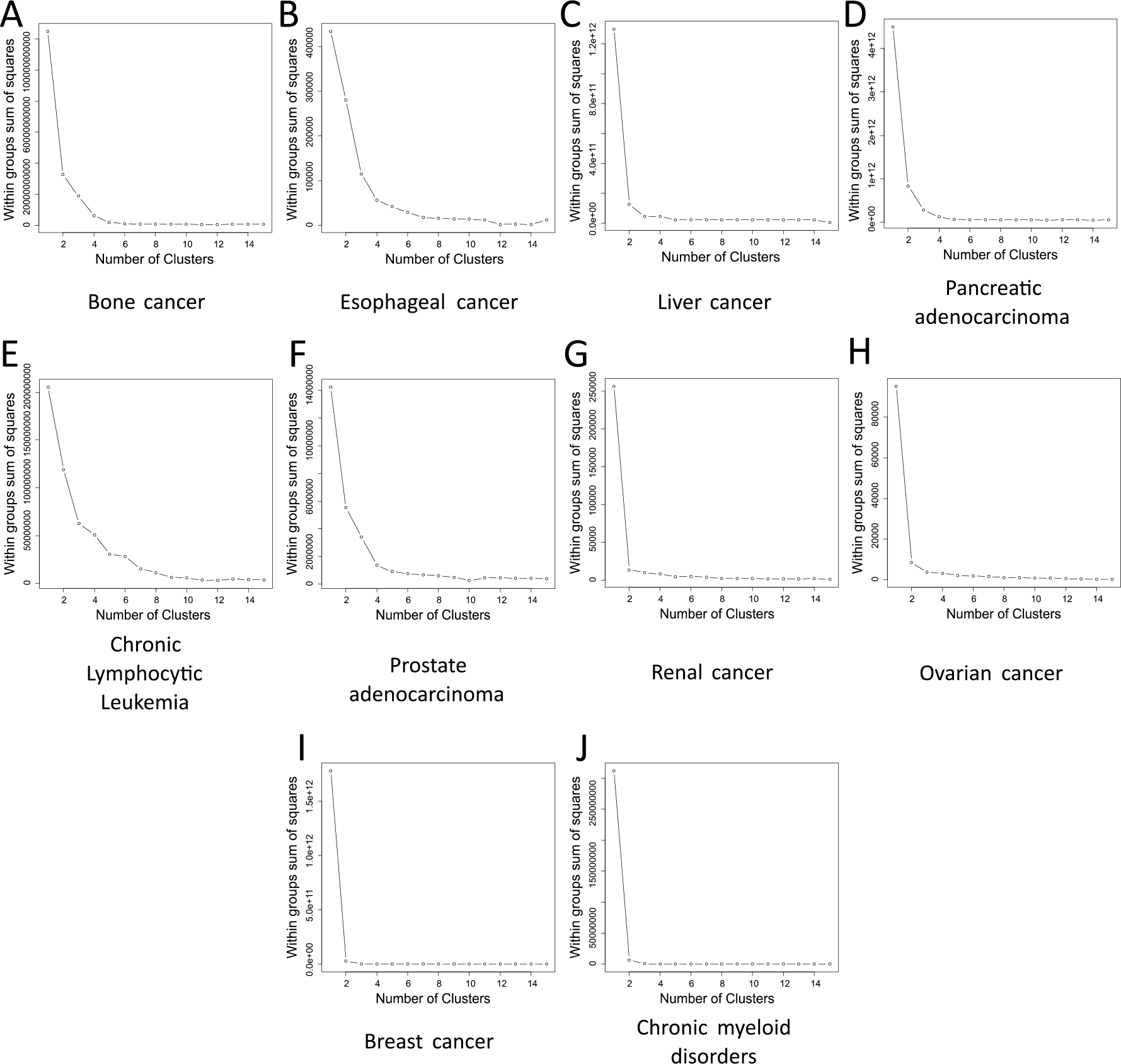
„Elbow method” to determine k for k-means clustering|. A-J, plot of reducing within group sum of squares for increasing k (number of clusters) in k-means clustering (Supplementary figure 4, Supplementary figure 5) of log_2_-transformed RPPB of all species-level taxa identified as tumor-linked and detected after filtering in all tumor-tissues for each indicated cancer.

**Supplementary Table 1:**
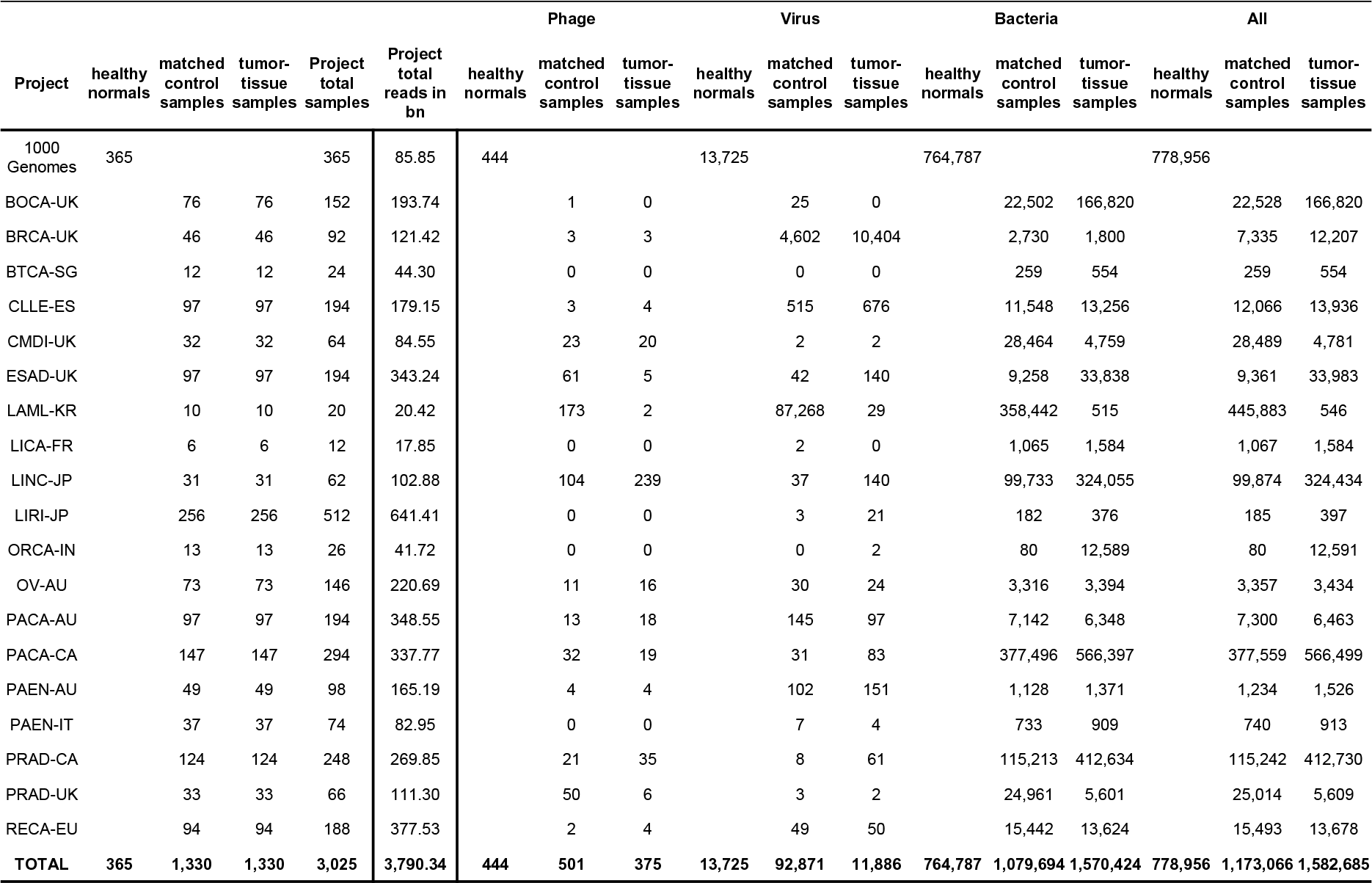
Included patients and samples

## REFERENCES

1. Schwabe, R. F. & Jobin, C. The microbiome and cancer. Nat. Rev. Cancer 13, 800–12 (2013).

2. Goodman, B. & Gardner, H. The microbiome and cancer. J. Pathol. 44, 667–676 (2018).

3. zur Hausen, H. Papillomaviruses and cancer: from basic studies to clinical application. Nat. Rev. Cancer 2, 342–350 (2002).

4. Tang, K.-W., Alaei-Mahabadi, B., Samuelsson, T., Lindh, M. & Larsson, E. The landscape of viral expression and host gene fusion and adaptation in human cancer. Nat. Commun. 4, 2513 (2013).

5. Cantalupo, P. G., Katz, J. P. & Pipas, J. M. Viral sequences in human cancer. Virology 513, 208–216 (2018).

6. Niedobitek, G. et al. Detection of human papillomavirus type 16 DNA in carcinomas of the palatine tonsil. J. Clin. Pathol. 43, 918–21 (1990).

7. Gillison, M. L. et al. Evidence for a Causal Association Between Human Papillomavirus and a Subset of Head and Neck Cancers. J. Natl. Cancer Inst. 92, 709–720 (2000).

8. Walboomers, J. M. M. et al. Human papillomavirus is a necessary cause of invasive cervical cancer worldwide. J. Pathol. 189, 12–19 (1999).

9. Schwarz, E. et al. Structure and transcription of human papillomavirus sequences in cervical carcinoma cells. Nature 314, 111–114 (1985).

10. Perz, J. F., Armstrong, G. L., Farrington, L. A., Hutin, Y. J. F. & Bell, B. P. The contributions of hepatitis B virus and hepatitis C virus infections to cirrhosis and primary liver cancer worldwide. J. Hepatol. 45, 529–538 (2006).

11. Shafritz, D. A., Shouval, D., Sherman, H. I., Hadziyannis, S. J. & Kew, M. C. Integration of Hepatitis B Virus DNA into the Genome of Liver Cells in Chronic Liver Disease and Hepatocellular Carcinoma. N. Engl. J. Med. 305, 1067–1073 (1981).

12. zur Hausen, H. The search for infectious causes of human cancers: Where and why. Virology 392, 1–10 (2009).

13. Uemura, N. et al. *Helicobacter pylori* Infection and the Development of Gastric Cancer. N. Engl. J. Med. 345, 784–789 (2001).

14. Watanabe, T., Tada, M., Nagai, H., Sasaki, S. & Nakao, M. Helicobacter pylori infection induces gastric cancer in mongolian gerbils. Gastroenterology 115, 642–8 (1998).

15. Castellarin, M. et al. Fusobacterium nucleatum infection is prevalent in human colorectal carcinoma. Genome Res. 22, 299–306 (2012).

16. Kostic, A. D. et al. Genomic analysis identifies association of Fusobacterium with colorectal carcinoma. Genome Res. 22, 292–8 (2012).

17. Repass, J. et al. Replication Study: Fusobacterium nucleatum infection is prevalent in human colorectal carcinoma. Elife 7, (2018).

18. Hanahan, D. & Weinberg, R. A. Hallmarks of cancer: the next generation. Cell 144, 646–74 (2011).

19. Goodwin, S., McPherson, J. D. & McCombie, W. R. Coming of age: ten years of next-generation sequencing technologies. Nat. Rev. Genet. 17, 333–351 (2016).

20. Riley, D. R. et al. Bacteria-Human Somatic Cell Lateral Gene Transfer Is Enriched in Cancer Samples. PLoS Comput. Biol. 9, e1003107 (2013).

21. Khoury, J. D. et al. Landscape of DNA virus associations across human malignant cancers: analysis of 3,775 cases using RNA-Seq. J. Virol. 87, 8916–26 (2013).

22. International Cancer Genome Consortium, T. I. C. G. et al. International network of cancer genome projects. Nature 464, 993–8 (2010).

23. Gibbs, R. A. et al. A global reference for human genetic variation. Nature 526, 68–74 (2015).

24. Wood, D. E. & Salzberg, S. L. Kraken: ultrafast metagenomic sequence classification using exact alignments. Genome Biol. 15, R46 (2014).

25. Li, H. & Durbin, R. Fast and accurate long-read alignment with Burrows-Wheeler transform. Bioinformatics 26, 589–95 (2010).

26. Caygill, C. P., Hill, M. J., Braddick, M. & Sharp, J. C. Cancer mortality in chronic typhoid and paratyphoid carriers. *Lancet (London*, England) 343, 83–4 (1994).

27. Scanu, T. et al. Salmonella Manipulation of Host Signaling Pathways Provokes Cellular Transformation Associated with Gallbladder Carcinoma. Cell Host Microbe 17, 763– 774 (2015).

28. Geller, L. T. et al. Potential role of intratumor bacteria in mediating tumor resistance to the chemotherapeutic drug gemcitabine. Science (80-.). 357, 1156–1160 (2017).

29. Feng, Y. et al. Metagenomic and metatranscriptomic analysis of human prostate microbiota from patients with prostate cancer. BMC Genomics 20, 146 (2019).

30. Kripalani-Joshi, S. & Law, H. Y. Identification of integrated epstein-barr virus in nasopharyngeal carcinoma using pulse field gel electrophoresis. Int. J. Cancer 56, 187– 192 (1994).

31. Morton, C. et al. Mapping of the human Blym-1 transforming gene activated in Burkitt lymphomas to chromosome 1. Science (80-.). 223, 173–175 (1984).

32. Robinson, K. M., Crabtree, J., Mattick, J. S. A., Anderson, K. E. & Dunning Hotopp, J. C. Distinguishing potential bacteria-tumor associations from contamination in a secondary data analysis of public cancer genome sequence data. Microbiome 5, 9 (2017).

33. Thompson, K. J. et al. A comprehensive analysis of breast cancer microbiota and host gene expression. PLoS One 12, e0188873 (2017).

34. Mazzoni, E. et al. Significant Association Between Human Osteosarcoma and Simian Virus 40. doi:10.1002/cncr.29137

35. Ramanan, P., Deziel, P. J. & Wengenack, N. L. Gordonia bacteremia. J. Clin. Microbiol. 51, 3443–7 (2013).

36. Ding, X. et al. Bacteremia due to Gordonia polyisoprenivorans: case report and review of literature. BMC Infect. Dis. 17, 419 (2017).

37. Gupta, M., Prasad, D., Khara, H. S. & Alcid, D. A rubber-degrading organism growing from a human body. Int. J. Infect. Dis. 14, e75–e76 (2010).

38. Zenz, T. et al. TP53 Mutation and Survival in Chronic Lymphocytic Leukemia. J. Clin. Oncol. 28, 4473–4479 (2010).

39. Austen, B. et al. Mutations in the ATM gene lead to impaired overall and treatment-free survival that is independent of IGVH mutation status in patients with B-CLL. (2005). doi:10.1182/blood-2004-11-4516

40. Balthazar, E. J., Megibow, A. J. & Hulnick, D. H. Cytomegalovirus esophagitis and gastritis in AIDS. AJR. Am. J. Roentgenol. 144, 1201–4 (1985).

41. Lepiller, Q., Tripathy, M. K., Di Martino, V., Kantelip, B. & Herbein, G. Increased HCMV seroprevalence in patients with hepatocellular carcinoma. Virol. J. 8, 485 (2011).

42. Leonardsson, H., Hreinsson, J. P., Löve, A. & Björnsson, E. S. Hepatitis due to Epstein– Barr virus and cytomegalovirus: clinical features and outcomes. Scand. J. Gastroenterol. 52, 893–897 (2017).

43. Bruix, J. & Llovet, J. M. Hepatitis B virus and hepatocellular carcinoma. J. Hepatol. 39, 59–63 (2003).

44. Lai, C.-C. et al. Infections caused by unusual Methylobacterium species. J. Clin. Microbiol. 49, 3329–31 (2011).

45. Langevin, S., Vincelette, J., Bekal, S. & Gaudreau, C. First case of invasive human infection caused by Cupriavidus metallidurans. J. Clin. Microbiol. 49, 744–5 (2011).

46. Ugge, H. et al. Acne in late adolescence and risk of prostate cancer. Int. J. cancer 142, 1580–1585 (2018).

47. Davidsson, S. et al. Frequency and typing of Propionibacterium acnes in prostate tissue obtained from men with and without prostate cancer. Infect. Agent. Cancer 11, 26 (2016).

48. Cohen, R. J., Shannon, B. A., Mcneal, J. E., Shannon, T. & Garrett, K. L. *Propionibacterium Acnes* Associated with Inflammation in Radical Prostatectomy Specimens: A Possible Link to Cancer Evolution? J. Urol. 173, 1969–1974 (2005).

49. Mahlen, S. D. Serratia infections: from military experiments to current practice. Clin. Microbiol. Rev. 24, 755–91 (2011).

50. Fessler, J., Matson, V. & Gajewski, T. F. Exploring the emerging role of the microbiome in cancer immunotherapy. J. Immunother. Cancer 7, 108 (2019).

51. Motzer, R. J. et al. Nivolumab versus Everolimus in Advanced Renal-Cell Carcinoma. N. Engl. J. Med. 373, 1803–13 (2015).

52. Sung, W.-K. et al. Genome-wide survey of recurrent HBV integration in hepatocellular carcinoma. Nat. Genet. Vol. 44, (2012).

53. Welcome | ICGC Data Portal. Available at: https://dcc.icgc.org/. (Accessed: 6th June 2019)

54. Index von /vol1/ftp/. Available at: ftp://ftp.1000genomes.ebi.ac.uk/vol1/ftp/. (Accessed: 6th June 2019)

55. European Nucleotide Archive & EMBL-EBI. Available at: https://www.ebi.ac.uk/ena. (Accessed: 6th June 2019)

56. Afgan, E. et al. The Galaxy platform for accessible, reproducible and collaborative biomedical analyses: 2018 update. Nucleic Acids Res. 46, W537–W544 (2018).

57. Li, H. et al. The Sequence Alignment/Map format and SAMtools. Bioinformatics 25, 2078–2079 (2009).

58. Picard Tools - By Broad Institute. Available at: http://broadinstitute.github.io/picard/. (Accessed: 6th June 2019)

59. Quinlan, A. R. & Hall, I. M. BEDTools: a flexible suite of utilities for comparing genomic features. Bioinformatics 26, 841–842 (2010).

60. Bolger, A. M., Lohse, M. & Usadel, B. Trimmomatic: a flexible trimmer for Illumina sequence data. Bioinformatics 30, 2114–20 (2014).

61. Blankenberg, D. et al. Manipulation of FASTQ data with Galaxy. Bioinformatics 26, 1783–5 (2010).

62. Rognes, T., Flouri, T., Nichols, B., Quince, C. & Mahé, F. VSEARCH: a versatile open source tool for metagenomics. PeerJ 4, e2584 (2016).

63. The R Foundation for Statistical Computing R version 3.3.2. (https://www.r-project.org/).

64. Escapa, I. F. et al. New Insights into Human Nostril Microbiome from the Expanded Human Oral Microbiome Database (eHOMD): a Resource for the Microbiome of the Human Aerodigestive Tract. mSystems 3, e00187–18 (2018).

65. Dewhirst, F. E. et al. The Human Oral Microbiome. J. Bacteriol. 192, 5002–5017 (2010).

66. HOMD :: Human Oral Microbiome Database. Available at: http://www.homd.org/index.php?name=HOMD. (Accessed: 6th June 2019)

67. de Goffau, M. C. et al. Recognizing the reagent microbiome. Nat. Microbiol. 3, 851– 853 (2018).

68. Salter, S. J. et al. Reagent and laboratory contamination can critically impact sequence-based microbiome analyses. BMC Biol. 12, 87 (2014).

69. Jervis-Bardy, J. et al. Deriving accurate microbiota profiles from human samples with low bacterial content through post-sequencing processing of Illumina MiSeq data. Microbiome 3, 19 (2015).

70. Leon, L. J. et al. Enrichment of Clinically Relevant Organisms in Spontaneous Preterm-Delivered Placentas and Reagent Contamination across All Clinical Groups in a Large Pregnancy Cohort in the United Kingdom. Appl. Environ. Microbiol. 84, e00483–18 (2018).

71. Laurence, M., Hatzis, C. & Brash, D. E. Common Contaminants in Next-Generation Sequencing That Hinder Discovery of Low-Abundance Microbes. PLoS One 9, e97876 (2014).

72. Glassing, A., Dowd, S. E., Galandiuk, S., Davis, B. & Chiodini, R. J. Inherent bacterial DNA contamination of extraction and sequencing reagents may affect interpretation of microbiota in low bacterial biomass samples. Gut Pathog. 8, 24 (2016).

73. Camacho, C., et al. BLAST+: architecture and applications. BMC Bioinformatics 10, 421 (2009).

74. Wally, N. et al. Plasmid DNA contaminant in molecular reagents. Sci. Rep. 9, 1652 (2019).

75. Embedding projector - visualization of high-dimensional data. Available at: https://projector.tensorflow.org/. (Accessed: 6th June 2019)

76. Morpheus. Available at: https://software.broadinstitute.org/morpheus/. (Accessed: 6th June 2019)

77. Li, H. & Durbin, R. Fast and accurate short read alignment with Burrows-Wheeler transform. Bioinformatics 25, 1754–1760 (2009).

78. Wick, R. R., Judd, L. M., Gorrie, C. L. & Holt, K. E. Unicycler: Resolving bacterial genome assemblies from short and long sequencing reads. PLOS Comput. Biol. 13, e1005595 (2017).

79. Sondka, Z. et al. The COSMIC Cancer Gene Census: describing genetic dysfunction across all human cancers. Nat. Rev. Cancer 18, 696–705 (2018).

80. Warden, C. D., Yuan, Y.-C. & Wu, X. Optimal Calculation of RNA-Seq Fold-Change Values. International Journal of Computational Bioinformatics and In Silico Modeling 2, (2013).

81. Ashburner, M. et al. Gene ontology: tool for the unification of biology. The Gene Ontology Consortium. Nat. Genet. 25, 25–9 (2000).

